# Integrative Genomic Analysis and Functional Studies Reveal GP5, GRN, MPO and MCAM as Causal Protein Biomarkers for Platelet Traits

**DOI:** 10.1101/854216

**Authors:** Dong Heon Lee, Chen Yao, Arunoday Bhan, Thorsten Schlaeger, Joshua Keefe, Benjamin AT Rodriguez, Shih-Jen Hwang, Ming-Huei Chen, Daniel Levy, Andrew D Johnson

## Abstract

**Rationale:** Mean platelet volume (MPV) and platelet count (PLT) are platelet measures that have been linked to cardiovascular disease (CVD) and mortality risk. Identifying protein biomarkers for these measures may yield insights into CVD mechanisms.

**Objective:** We aimed to identify causal protein biomarkers for MPV and PLT among 71 CVD-related plasma proteins measured in Framingham Heart Study (FHS) participants.

**Methods and Results:** We conducted integrative analyses of genetic variants associated with PLT and MPV with protein quantitative trait locus (pQTL) variants associated with plasma proteins followed by Mendelian randomization (MR) to infer causal relations of proteins for PLT/MPV, and tested protein-PLT/MPV association in FHS participants. Utilizing induced pluripotent stem cell (iPSC)-derived megakaryocyte (MK) clones that produce functional platelets, we conducted RNA-sequencing and analyzed transcriptome-wide differences between low- and high-platelet producing clones. We then performed small interfering RNA (siRNA) gene knockdown experiments targeting genes encoding proteins with putatively causal platelet effects in MK clones to examine effects on platelet production. Protein-trait association analyses were conducted for MPV (n = 4,348) and PLT (n = 4,272). Eleven proteins were associated with MPV and 31 with PLT. MR identified four putatively causal proteins for MPV and four for PLT. Glycoprotein V (GP5), granulin (GRN), and melanoma cell adhesion molecule (MCAM) were associated with PLT in both protein-trait and MR analyses. Myeloperoxidase (MPO) showed significant association with MPV in both analyses. MK RNA-sequencing analysis results were directionally concordant with observed and MR-inferred associations for GP5, GRN, and MCAM. In siRNA gene knockdown experiments, silencing GP5, GRN, and MPO decreased platelet counts.

**Conclusions:** By integrating population genomics data, epidemiological data, and iPSC-derived MK experiments, we identified four proteins that are causally linked to platelet counts. These proteins and genes may be further explored for their utility in increasing platelet production in bioreactors for transfusion medicine purposes as well as their roles in the pathogenesis of CVD via a platelet/blood coagulation-based mechanism.

## Introduction

Platelets are circulating anucleate cells produced by megakaryocytes (MK) in the bone marrow^1^ that initiate thrombus formation at the site of injury^2^. They also function as innate and adaptive immunity regulators through secretion of antibacterial proteins, interactions with leukocytes, and mediation of pro-inflammatory functions of neutrophils and dendritic cells^3^. Their central roles in thrombus formation and immune response make platelets notable contributors to the pathobiology of atherosclerosis^4^ and arterial and venous thrombosis^5^, including venous thromboembolism (VTE)^6^. When the endothelium of a vessel is damaged, platelets aggregate at the site of injury and can promote atherosclerosis and atherothrombosis^5^, which in turn can cause acute coronary syndromes and strokes^7^. A widely-used platelet clinical metric, mean platelet volume (MPV)^8^, has been reported to be associated with risk of acute myocardial infarction (AMI) events^9^, coronary artery disease (CAD)^10^, and VTE^6^. Prior studies have also shown that higher MPV is associated with greater mortality risk among AMI patients^9^ and patients with sepsis^11^. Furthermore, greater MPV is also correlated with poor prognosis for patients with infective endocarditis^12^ and risk of pulmonary embolism among patients with deep vein thrombosis^13^. A prospective cohort study by Mayer and colleagues demonstrated that elevated MPV levels were associated with increased risk of major adverse cardiovascular events among patients with asymptomatic carotid atherosclerosis^14^. Platelet count (PLT), on the other hand, has a U-shaped association with mortality; high- and low-platelet counts have been reported to be associated with increased risk of coronary heart disease (CHD) and cancer mortality^15^ and with overall mortality in people older than 65 years of age^16, 17^.Therefore, given the clinical importance of platelets vis-à-vis cardiovascular diseases (CVD), identifying putatively causal protein biomarkers associated with MPV and PLT may lead to a better understanding of disease mechanisms and highlight therapeutic targets for the treatment and prevention of CVD.

As part of the Systems Approach to Biomarker Research in Cardiovascular Disease (SABRe CVD) Initiative, we conducted a genome-wide association study (GWAS) to identify genetic loci associated with circulating levels of 71 CVD-related plasma proteins (pQTLs; protein quantitative trait loci) in 7333 Framingham Heart Study (FHS) participants^18^. In the present study we sought to identify associations between circulating protein levels and platelet phenotypes.

Identifying overlap between genetic variants associated with circulating protein levels and single nucleotide polymorphisms (SNPs) associated with complex traits from previously published genome-wide association studies (GWAS) can prioritize candidate protein biomarkers that can be further explored as causal proteins for disease and/or drug targets of complex disease phenotypes. We previously demonstrated the utility of this technique by identifying putatively causal protein biomarkers for CVD^18^ and chronic obstructive pulmonary disease^19^. Because PLT and MPV are highly heritable quantitative traits^20^ with large published GWAS and exome studies^21–23^, we hypothesized that these traits may be amenable to a similar approach to identify causal protein biomarkers. To that end, we conducted Mendelian randomization (MR) analyses to test for putatively causal associations between protein biomarkers and MPV and PLT. Proteins that were associated with MPV or PLT in both FHS data and MR were functionally validated using RNA-sequencing data of human induced pluripotent stem cells (iPSC) MK clones followed by siRNA gene knockdown experiments.

## Methods

Study Design: An overall flowchart of our study design is shown in Figure 1. We first conducted protein-trait association analyses using FHS data^24^ for MPV and PLT. We then performed MR using *cis-*pQTL variants^18^ that overlapped with prior MPV/ PLT GWAS^21, 22^ SNPs as instrumental variables to test for putatively causal protein-MPV/PLT associations. Proteins showing significant causal association were further investigated for transcriptome-level association with platelet production using RNA sequencing data from human iPSC MK. Finally, we performed gene knockdown experiments in iPSC MK.

**Figure 1.**
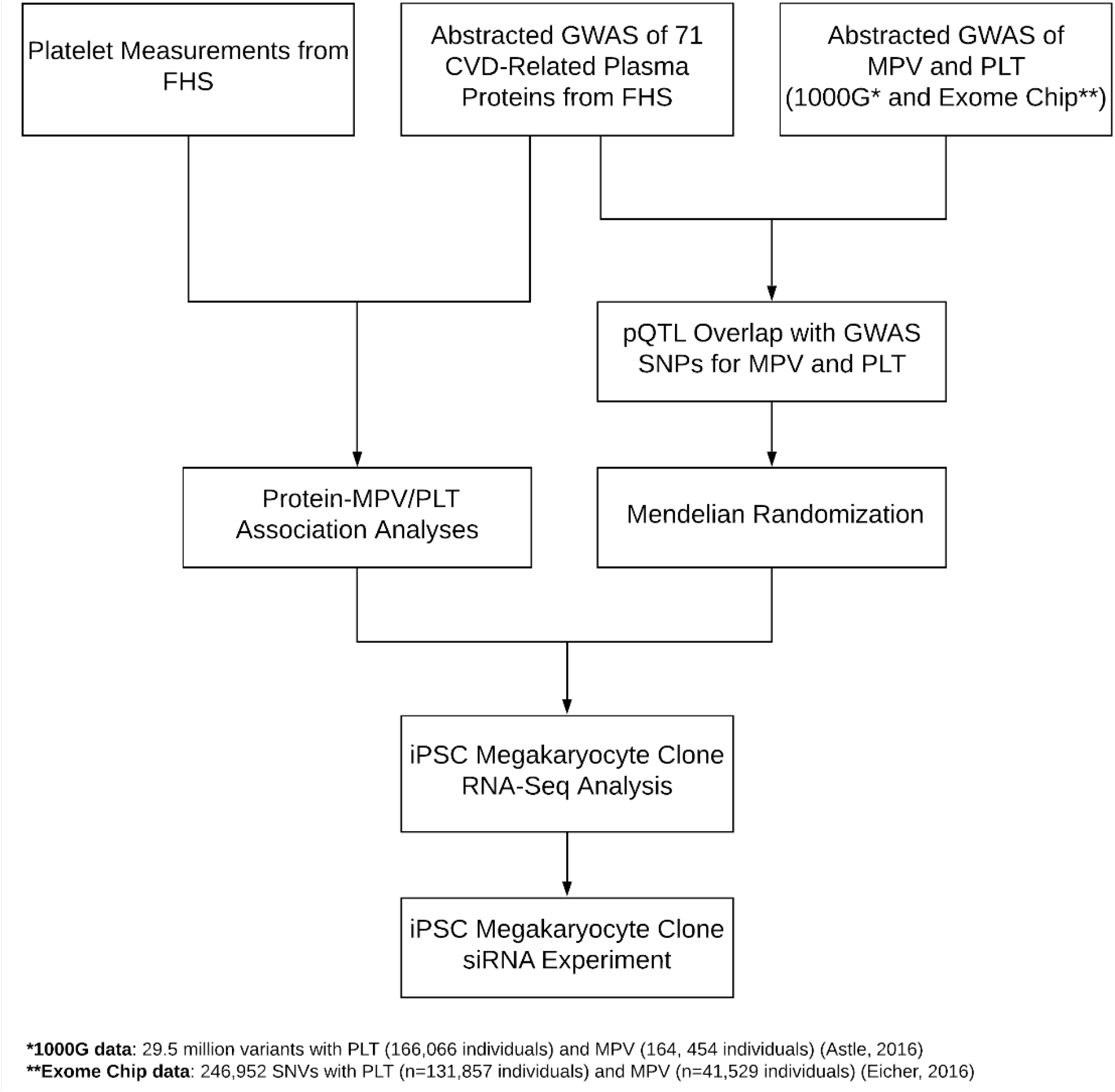
Flowchart of Study Design.

### Protein-Trait Association Analysis

Study participants: The study sample consisted of FHS Offspring and Third Generation cohort participants. The FHS Offspring study was comprised of 5124 children or spouses of original FHS cohort participants; enrollment began in 1971^25, 26^. The Third Generation cohort included 4095 children of Offspring cohort participants who began enrollment in 2002^27^. Among the Third Generation cohort participants, 3411 attended Examination 2 (2008-2011) and provided blood samples for the measurements of PLT and MPV. Among the Offspring cohort participants, 2430 attended the 9^th^ Examination (2011-2014) and provided information on cardiovascular health and blood samples for laboratory tests including PLT and MPV. Of these 5841 Offspring and Third Generation participants with platelet measurements, 5233 provided informed consent and participated in the Systems Approach to Biomarker Research in Cardiovascular Disease (SABRe CVD) Initiative^24^ for which plasma protein measurements were measured. Plasma proteins were measured in plasma samples obtained at Examination 7 (1998-2001) for the Offspring cohort participants and at Examination 1 (2002-2005) for the Third Generation cohort participants. Among these 5233 participants, complete information on PLT, MPV, and normalized plasma protein concentration was available in 4349 participants including 2026 Offspring (46.6%) and 2323 Third Generation participants (53.4%). Characteristics of 4348 participants with MPV measurements and 4272 participants with PLT measurements are summarized in **Table 1**.

**Table 1.** Characteristics of FHS Participants with Platelet Measurements.

|  | **MPV (N = 4,348)** |  | **PLT (N = 4,272)** |  |
| --- | --- | --- | --- | --- |
| **Characteristic** | **Mean** | **SD** | **Mean** | **SD** |
| Age (year) | 58.1 | 14.5 | 57.9 | 14.4 |
| BMI (kg/m<sup>2</sup>) | 28.3 | 5.6 | 28.3 | 5.6 |
| Mean Platelet Volume (fL) | 8.7 | 0.96 | - | - |
| Platelet Count (10<sup>3</sup> cells/μL) | - | - | 240.7 | 58.7 |
| Systolic BP (mmHg) | 121.5 | 16.0 | 121.4 | 15.9 |
| Diastolic BP (mmHg) | 73.3 | 9.5 | 73.4 | 9.5 |
| Total-cholesterol (mg/dL) | 185.5 | 36.1 | 185.8 | 36.0 |
| HDL-cholesterol (mg/dL) | 60.7 | 18.6 | 60.9 | 18.7 |
| Triglyceride (mg/dL) | 115.2 | 70.0 | 115.0 | 70.1 |
|  | **N** | **%** | **N** | **%** |
| Women | 2276 | 52.4 | 2256 | 52.8 |
| Cigarette Smoker | 361 | 8.3 | 357 | 8.4 |
| Diabetic | 329 | 7.7 | 321 | 7.6 |
| Hypertension Treatment | 1560 | 36.0 | 1513 | 35.6 |
| Hyperlipidemia Treatment | 1425 | 32.8 | 1383 | 32.4 |
| Anticoagulant Use | 307 | 7.1 | 294 | 6.9 |
| Prevalent CVD | 392 | 9.0 | 371 | 8.7 |

Hematological measures: Blood PLT and MPV were measured using the Coulter HmX hematology analyzer (Beckman Coulter, Inc.)^8^.

Protein quantification: All plasma proteins were measured as part of the Systems Approach to Biomarker Research in Cardiovascular Disease (SABRe CVD) Initiative^24^. The protein quantification procedure has been described previously^18^. Briefly, fasting blood plasma samples were analyzed using a modified enzyme-linked immunosorbent assay sandwich method, multiplexed on Luminex xMAP platform (Luminex, Inc., Austin, TX), for plasma protein levels^18^.

Statistical methods: We applied linear mixed effects models (LME) to characterize significant associations between 71 plasma proteins and platelet measurements (MPV and PLT). We adjusted for sex, age at the time of platelet measurement, age squared, and ten genotype-based principal components, and derived the residuals. The residuals were inverse normalized and tested for association with the 71 plasma proteins using LME. Bonferroni corrected significance threshold was used to identify significant biomarkers associated with platelet measurements.

### Causal genetic polymorphisms

Identifying overlap of pQTL variants with GWAS variants: We interrogated our pQTL database^18^ (n=6861 participants were included in the GWAS sample with 1000 Genomes Project reference panel (1000G build 37 phase 1 v3) imputed dosage data) for overlap of *cis*-pQTL and *trans-*pQTL variants with single nucleotide polymorphisms (SNPs) from prior genome-wide association studies (GWAS) of MPV and PLT^21, 22^. Only *cis-*pQTLs that passed Bonferroni-corrected (BF) significance threshold at P < 1.25E-07 and *trans-*pQTLs that passed BF significance threshold at P < 7.04E-10 were considered for overlap analysis^18^.

MR testing for causal protein-trait association: MR was conducted using the TwoSampleMR^28^ R package using pruned *cis-*pQTL variants (linkage disequilibrium [LD] threshold of r^2^ < 0.1) as instrumental variables. Among the 71 CVD-related proteins, 37 proteins had *cis*-pQTL variants that were suitable as instrumental variables in MR. The Wald ratio, which is a ratio of the regression coefficient of the exposure on the instrumental variable to that of the outcome on the instrumental variable^29^, was used for proteins with one independent *cis*-pQTL, and inverse-variance weighted regression was used for proteins with multiple independent *cis*-pQTL variants^30^.

### RNA sequencing analysis

RNA sequencing analysis and derived platelet counting from human induced pluripotent stem cell (iPSC) derived immortalized megakaryocytic cell line (imMKCL) derived subclones: Expansion of imMKCLs: ImMKCLs are cultured and expanded via doxycycline dependent expression of C-MYC, BMI-1 and BCL-XL^31^. ImMKCLs are maintained in a humidified incubator at 37°C and 5% CO2 and in IMDM (Sigma-Aldrich) medium supplemented with 15% fetal bovine serum (FBS; Sigma-Aldrich), L-glutamine (Gibco), Insulin-transferrin-selenium (Gibco), 50 mg/mL Ascorbic acid (A4544; Sigma-Aldrich), and 450 mM 1-Thioglycerol (Sigma-Aldrich), 50 ng/mL carrier-free recombinant human stem cell factor (SCF; R&D Systems), 50 ng/mL TPO (R&D Systems), and 5 mg/mL Doxycycline (Clontech).

Differentiation of imMKCLs to generate platelets: Induction of imMKCL maturation was carried out in a humidified incubator at 37C and 5% CO2 with for 6 days in IMDM medium supplemented with 15% FBS consisted of 50 ng/mL SCF, 50 ng/mL TPO, 15 mM KP-457 (Medchemexpress), 0.5 mM GNF-351 (TOCRIS), and 10 mM Y27632 (Medchemexpress). On day 3 imMKCLs are highly polyploid and poised for platelet generation and platelet generation occurs between day 5 and 6.

Since heterogenous (Het) imMKCLs consisted of both non-platelet producing and platelet producing megakaryocytes, the imMKCLs were subcloned to decrease the ratio of non-platelet producing and platelet producing megakaryocytes. Subcloning of imMKCLs resulted in derivation of functionally distinct subclones. The 7 subclones varied in their ability to generate platelets and were labeled as Clone 42, Clone JC, Clone 32, Clone DKO, Clone 38, Clone 62, and Clone 72. To identify the genes and signaling networks that are necessary for platelet generation, we performed transcriptomic analyses of the 8 clones (Het and 7 subclones). Three biological replicates for each of the 8 clones were created, and maturing imMKCLs were harvested on day 3 for RNA extraction and subsequent platelet output measurements of the respective subclones were performed on day 6 via flow cytometry. Total RNA was extracted using miRNeasy Mini Kit (QIAGEN) according to the manufacturer’s instructions. RNA-seq libraries were prepared using the NEB Ultra (PolyA) kit as per manufacturer’s protocol with 50 ng input RNA. RNA-seq libraries were prepared and sequenced using the 200-cycle paired-end kit on the Illumina HiSeq2500 system. RNA-seq reads were analyzed with the Tuxedo Tools following a standard protocol. Reads were mapped with TopHat version 2.1.0 and Bowtie2 version 2.2.4 with default parameters against build hg19 of the human genome, and build hg19 of the RefSeq human genome annotation. Then, association analysis between gene transcript levels (fpkm) and platelet output (number of platelets per megakaryocyte) was performed via linear mixed effects model to account for the eight clone types.

### siRNA gene knockdown experiments

Immortalized megakaryocyte progenitor cell lines (imMKCL) culture: imMKCLs were generated from human induced pluripotent stem cells (iPSCs) and maintained in presence of 5 μg/mL doxycycline (DOX) as previously described^31^. Removal of DOX results in imMKCL maturation and the generation of platelets after six days.

siRNA transfection assay: The imMKCLs growing in the differentiation medium (without DOX) on day 4 were seeded in a 24-well plate, 24 hours prior to transfection. The cells were transfected with 2ug esiRNA (mixture of siRNA oligos, (Sigma-aldrich)) specific to MCAM, MPO and GRN as well as 500nM of siRNA targeted against GP5 or control eGFP siRNA oligonucleotides using Lipofectamine 2000 (Thermo Fisher Scientific) according to the manufacturer’s instructions. 48 hours post-transfection, the cells were harvested for RT-qPCR and *in vitro* flow cytometric analysis of platelets.

RNA extraction, reverse transcription, and RT-qPCR: RNA extraction was performed using an RNAeasy kit (Qiagen). Reverse transcription was performed using Superscript III (Invitrogen), using Oligo (dT) 15 primer. Quantitative PCR was performed in triplicate with SYBR Green and CFX96 real-time PCR detection system (Bio-rad). Target transcript abundance was calculated relative to GAPDH (reference gene) using the 2-ΔΔCT method. Gene specific primer pairs are present in **Supplementary Table 1.** Flow cytometric analysis of platelet cell markers: Flow cytometric based platelet count analyses was performed on MCAM, MPO, GRN, GP5 or control GFP knockdown experiments on Day 6 imMKCLs grown in differentiation medium without DOX. Briefly, cell aliquots were incubated for 20 min with fluorescently labeled monoclonal antibodies and dyes. The antibodies used were: phycoerythrin (PE)- mouse anti-human anti–CD41a antibody (BD Pharmingen; Cat# 555467); fluorescein isothiocyanate (FITC)- mouse anti-human Annexin-V (biolegend; Cat# 640945) and APC-anti-human anti-CD42b (GPIb) monoclonal antibody (BD Pharmingen; Cat# 551061). The calcein AM blue was used to identify live platelet sized events. Stains were fixed in 0.5% paraformaldehyde solution. Data were collected on a LSRII flow cytometer (BD Biosciences) and analyzed with FlowJo software (FlowJo LLC). Calcein AM blue+ AnnexinV-CD41+CD42b+ events were determined as platelets.

## Results

### FHS protein-MPV/PLT association analyses

Among the 71 CVD-related plasma proteins, ten (14.1%) were associated with MPV after Bonferroni (BF) correction (P < 0.05/71) and another 17 (23.9%) proteins were nominally significant (P < 0.05). For PLT, 31 (43.7%) proteins showed significant association after BF correction and 13 (18.3%) were nominally significant (P < 0.05) (**Table 2** and **Supplementary Table 2**). Seven proteins showed significant association with both PLT and MPV: soluble CD40 ligand (sCD40L), stromal cell-derived factor 1 (SDF1), pro-platelet basic protein (PPBP), translocation associated notch homolog (Notch1), adrenomedullin (ADM), glyceraldehyde 3-phosphate dehydrogenase (GAPDH), and cadherin-13 (CDH13). All seven proteins’ direction of effect for MPV was the opposite of their direction of effect for PLT (Pearson’s correlation: −0.90, 95% CI: (−0.99, −0.48)), reflecting the inverse correlation of MPV and PLT^32^.

**Table 2.** Protein-MPV/PLT Association Analysis Results from FHS Data.

| Trait | Protein<sup>†</sup> | $\beta^*$ | SE | p-value** |
| --- | --- | --- | --- | --- |
| MPV | sCD40L<sup>††</sup> | -0.108 | 0.014 | 5.81E-14 |
|  | SDF1<sup>††</sup> | -0.108 | 0.015 | 2.47E-13 |
|  | PPBP<sup>††</sup> | -0.087 | 0.015 | 2.68E-09 |
|  | NOTCH1<sup>††</sup> | 0.066 | 0.015 | 7.71E-06 |
|  | ADM<sup>††</sup> | 0.066 | 0.015 | 9.74E-06 |
|  | GAPDH<sup>††</sup> | -0.064 | 0.014 | 9.74E-06 |
|  | MMP9 | 0.061 | 0.015 | 3.09E-05 |
|  | RETN | 0.063 | 0.015 | 4.71E-05 |
|  | MPO | 0.058 | 0.016 | 2.06E-04 |
|  | CDH13<sup>††</sup> | -0.050 | 0.015 | 5.63E-04 |
| PLT | PPBP<sup>††</sup> | 0.217 | 0.0150 | 2.80E-47 |
|  | sCD40L<sup>††</sup> | 0.183 | 0.0150 | 2.99E-34 |
|  | PAI1 | 0.157 | 0.0151 | 2.09E-25 |
|  | GMP140 | 0.152 | 0.0155 | 9.78E-23 |
|  | sRAGE | -0.136 | 0.0158 | 8.42E-18 |
|  | GP5 | 0.128 | 0.0155 | 1.31E-16 |
|  | CNTN1 | -0.122 | 0.0155 | 2.96E-15 |
|  | CDH13<sup>††</sup> | 0.118 | 0.0152 | 6.95E-15 |
|  | LDLR | 0.118 | 0.0152 | 9.69E-15 |
|  | GAPDH<sup>††</sup> | 0.101 | 0.0152 | 2.96E-11 |
|  | NRCAM | 0.0875 | 0.0150 | 5.87E-09 |
|  | PMP2 | 0.0878 | 0.0152 | 8.24E-09 |
|  | NOTCH1<sup>††</sup> | -0.0882 | 0.0155 | 1.17E-08 |
|  | SDF1<sup>††</sup> | 0.0848 | 0.0156 | 5.33E-08 |
|  | KLKB1 | 0.0849 | 0.0157 | 6.72E-08 |
|  | CRP | 0.0813 | 0.0151 | 6.82E-08 |
|  | ADAM15 | 0.0781 | 0.0151 | 2.40E-07 |
|  | CD56 (NCAM) | -0.0779 | 0.0156 | 6.50E-07 |
|  | ANGPTL3 | 0.0743 | 0.0152 | 1.09E-06 |
|  | HPX | 0.0742 | 0.0153 | 1.29E-06 |
|  | ApoB | 0.0731 | 0.0154 | 2.09E-06 |
|  | C2 | 0.0731 | 0.0154 | 2.19E-06 |
|  | AGP1 | 0.0713 | 0.0152 | 2.84E-06 |
|  | MCAM | -0.0715 | 0.0156 | 4.37E-06 |
|  | CD5L | -0.0640 | 0.0160 | 6.44E-05 |
|  | ADM<sup>††</sup> | -0.0615 | 0.0155 | 7.51E-05 |
|  | BGLAP | -0.0613 | 0.0155 | 7.65E-05 |
|  | FGG | 0.0617 | 0.0158 | 9.30E-05 |
|  | BCHE | 0.0592 | 0.0156 | 1.53E-04 |
|  | NTproBNP | -0.0591 | 0.0157 | 1.70E-04 |
|  | A1M | 0.0589 | 0.0161 | 2.50E-04 |

<sup>†</sup>A full list of proteins showing nominal significance ( $p < 0.05$ ) is in Supplementary Table 2
<sup>††</sup>Proteins showing significant associations with both MPV and PLT
\* Unit: per standard deviation increment in rank-based inverse normal transformed protein level
\*\*Bonferroni-corrected significance threshold for 71 proteins $P = 0.05/71 = 7.04E-04$

### pQTL variants that overlap with published GWAS of MPV and PLT

We identified 15 pQTLs (four *cis-* and 11 *trans-*pQTLs) corresponding to 15 proteins that overlapped with MPV-GWAS SNPs (**Table 3**). Among the 15 proteins, CDH13 was significantly associated with MPV in FHS data after BF correction (**Table 2**), and six proteins (sRAGE, PMP2, LDLR, ADAM15, CD14, and NRCAM) showed nominal significance. Seventeen pQTLS (five *cis-* and 12 *trans-*pQTLs) corresponding to 17 proteins overlapped with PLT-GWAS SNPs (**Table 4**). Among these 17 proteins, 12 proteins (GMP140, sRAGE, GP5, CDH13, LDLR, NRCAM, PMP2, KLKB1, ADAM15, ANGPTL3, C2, MCAM) passed BF correction threshold and were associated with PLT in FHS data (**Table 2**), and two proteins (CD14, GRN) showed nominal significance.

**Table 3.**
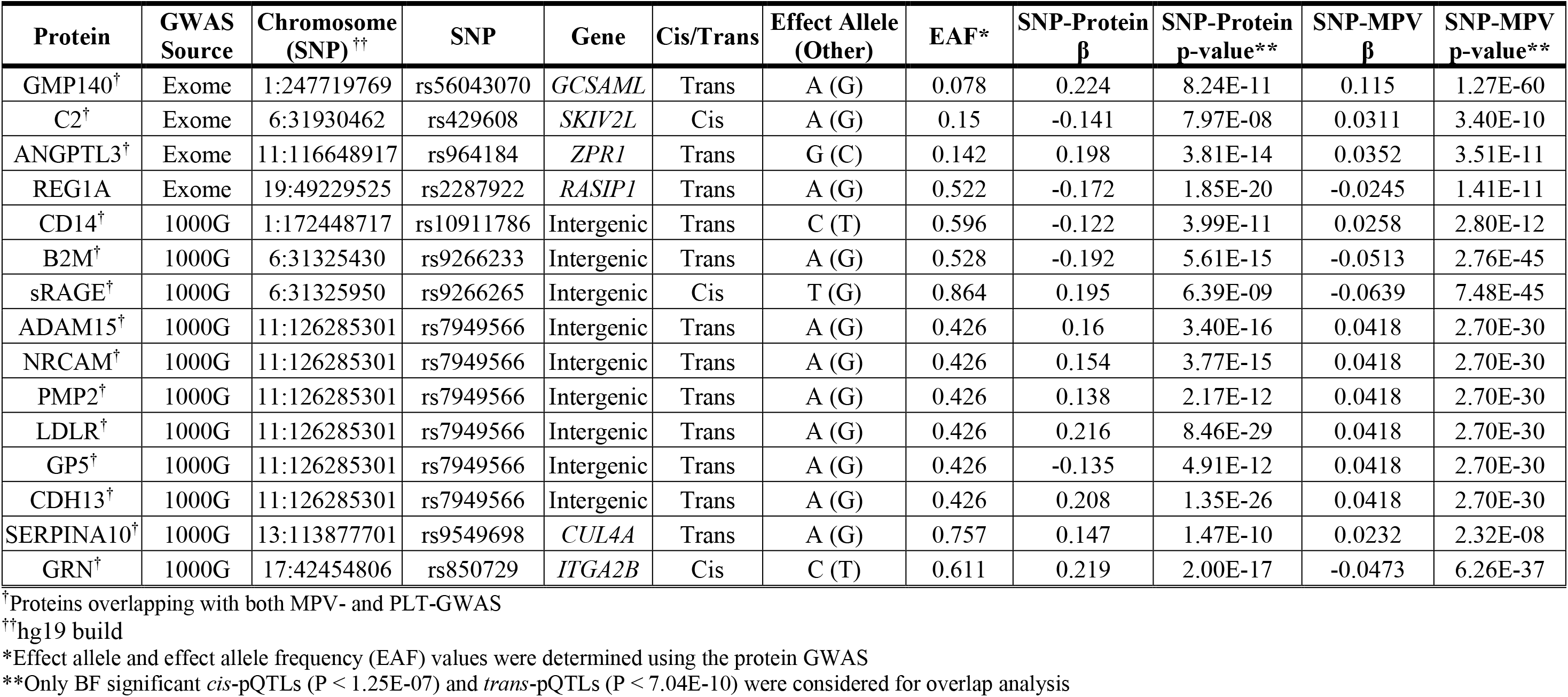
Overlap of pQTLs with MPV GWAS SNPs.

**Table 4.**
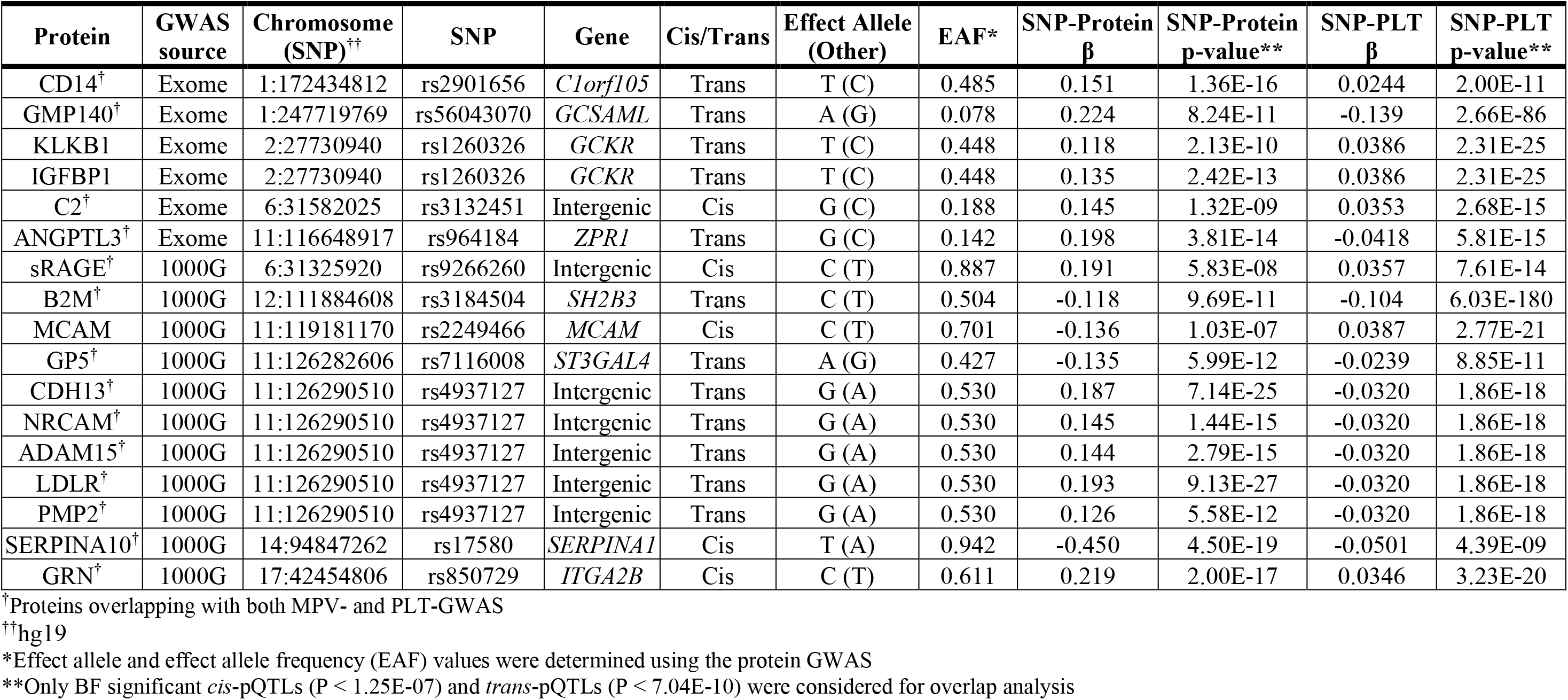
Overlap of pQTLs with PLT GWAS SNPs.

### MR analyses

We conducted MR analyses using 40 proteins with *cis*-pQTL variants. Three of the 40 proteins did not have pQTLs that overlapped with GWAS variants for MPV/PLT. Thus, 37 proteins were tested for putatively causal associations with MPV/PLT. MR analysis revealed four proteins that were causally associated with MPV after BF correction (P < 0.05/37) and five proteins that were nominally significant at P < 0.05 (**Table 5** and **Supplementary Table 3**). Similarly, four proteins were causally associated with PLT after BF correction and five showed nominally significant association (**Table 5** and **Supplementary Table 3**). SERPINA10 was tested for horizontal pleiotropy and heterogeneity because it had 28 SNPs (**Supplementary Table 4**); there was no evidence of horizontal pleiotropy (Egger regression intercept: 0.005, p-value: 0.14), but a forest plot of individual SNP MR effect size and a test based on the Egger method confirmed heterogeneity (p-value: 0.022) (**Supplementary Figure 1-2**). Removal of seven SERPINA10 *cis-*pQTL SNPs with negative MR coefficients resulted in modest improvement of the effect size and significance (IVW MR β: 0.019, SE: 0.0036, p-value: 1.69E-07) (**Supplementary Figure 3**). Granulin (GRN) and platelet glycoprotein V (GP5) were causally associated with both MPV and PLT (**Table 5**).

**Table 5.**
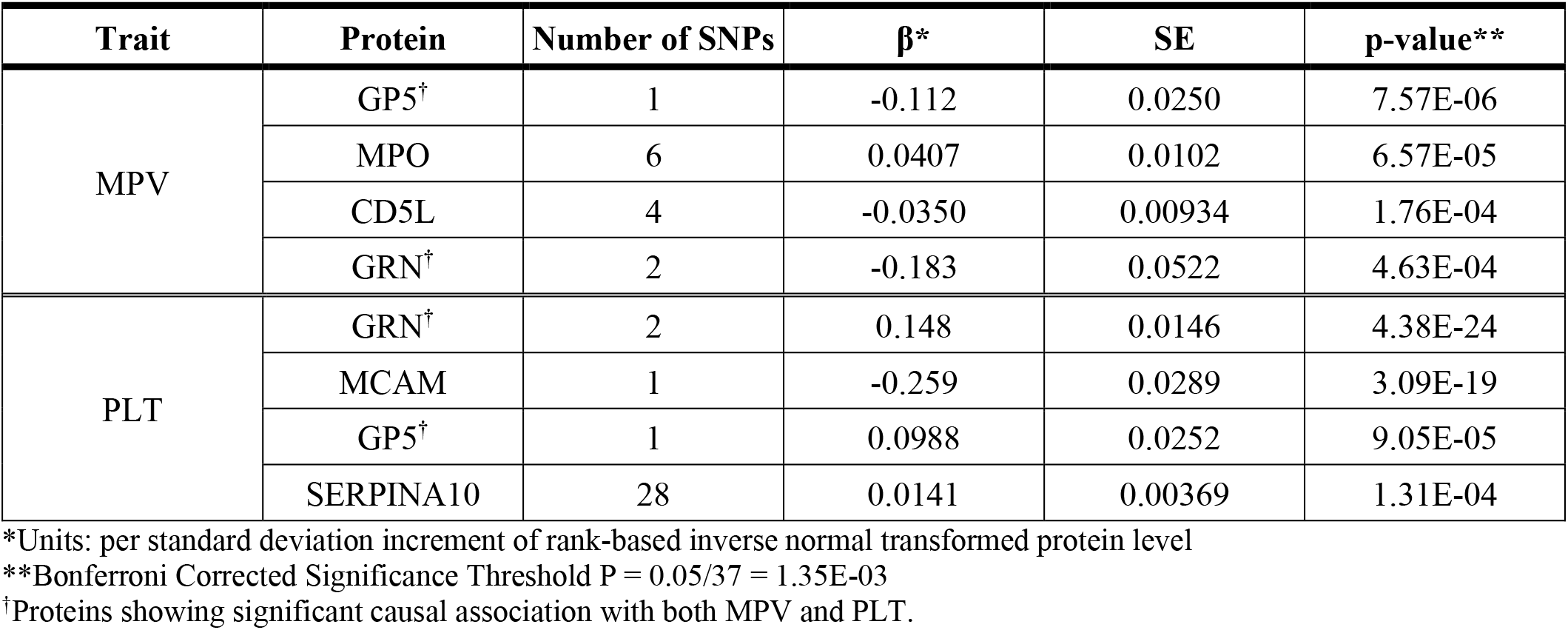
MR Results for MPV and PLT.

| Trait | Protein | Number of SNPs | $\beta^*$ | SE | p-value** |
| --- | --- | --- | --- | --- | --- |
| MPV | GP5<sup>†</sup> | 1 | -0.112 | 0.0250 | 7.57E-06 |
|  | MPO | 6 | 0.0407 | 0.0102 | 6.57E-05 |
|  | CD5L | 4 | -0.0350 | 0.00934 | 1.76E-04 |
|  | GRN<sup>†</sup> | 2 | -0.183 | 0.0522 | 4.63E-04 |
| PLT | GRN<sup>†</sup> | 2 | 0.148 | 0.0146 | 4.38E-24 |
|  | MCAM | 1 | -0.259 | 0.0289 | 3.09E-19 |
|  | GP5<sup>†</sup> | 1 | 0.0988 | 0.0252 | 9.05E-05 |
|  | SERPINA10 | 28 | 0.0141 | 0.00369 | 1.31E-04 |

\*Units: per standard deviation increment of rank-based inverse normal transformed protein level
\*\*Bonferroni Corrected Significance Threshold $P = 0.05/37 = 1.35E-03$
<sup>†</sup>Proteins showing significant causal association with both MPV and PLT.

Myeloperoxidase (MPO) was positively associated with MPV in both MR and protein-trait association analyses after BF correction, with directional consistency of effect estimates between the two analyses (Figure 2). N-terminal prohormone of brain natriuretic peptide (NTproBNP) was nominally significant in both analyses with directional consistency (Figure 2). Melanoma cell adhesion molecule (MCAM/CD146/MUC18) and glycoprotein V (GP5) were associated with PLT in both MR and protein-trait association analyses with directional consistency (Figure 2). GRN was BF significant in MR but nominally significant in protein-PLT analysis with directional consistency. Adrenomedullin (ADM), contactin 1 (CNTN1), and C-reactive protein (CRP) were BF significant in protein-PLT association analysis but nominally significant in MR; ADM showed directionally consistent effect estimates, but CNTN1 and CRP showed directional inconsistency (Figure 2). Instrumental variable SNPs for GP5, GRN, MCAM, and MPO (**Supplementary Table 4**), the genes carried forward in experiments, were queried for CHD outcome associations in CARDIoGRAMplusC4D^33^, UK BioBank GWAS^34^, the MEGASTROKE Consortium^35^, and the Japan BioBank^36^ (**Supplementary Table 5**).

**Figure 2.**
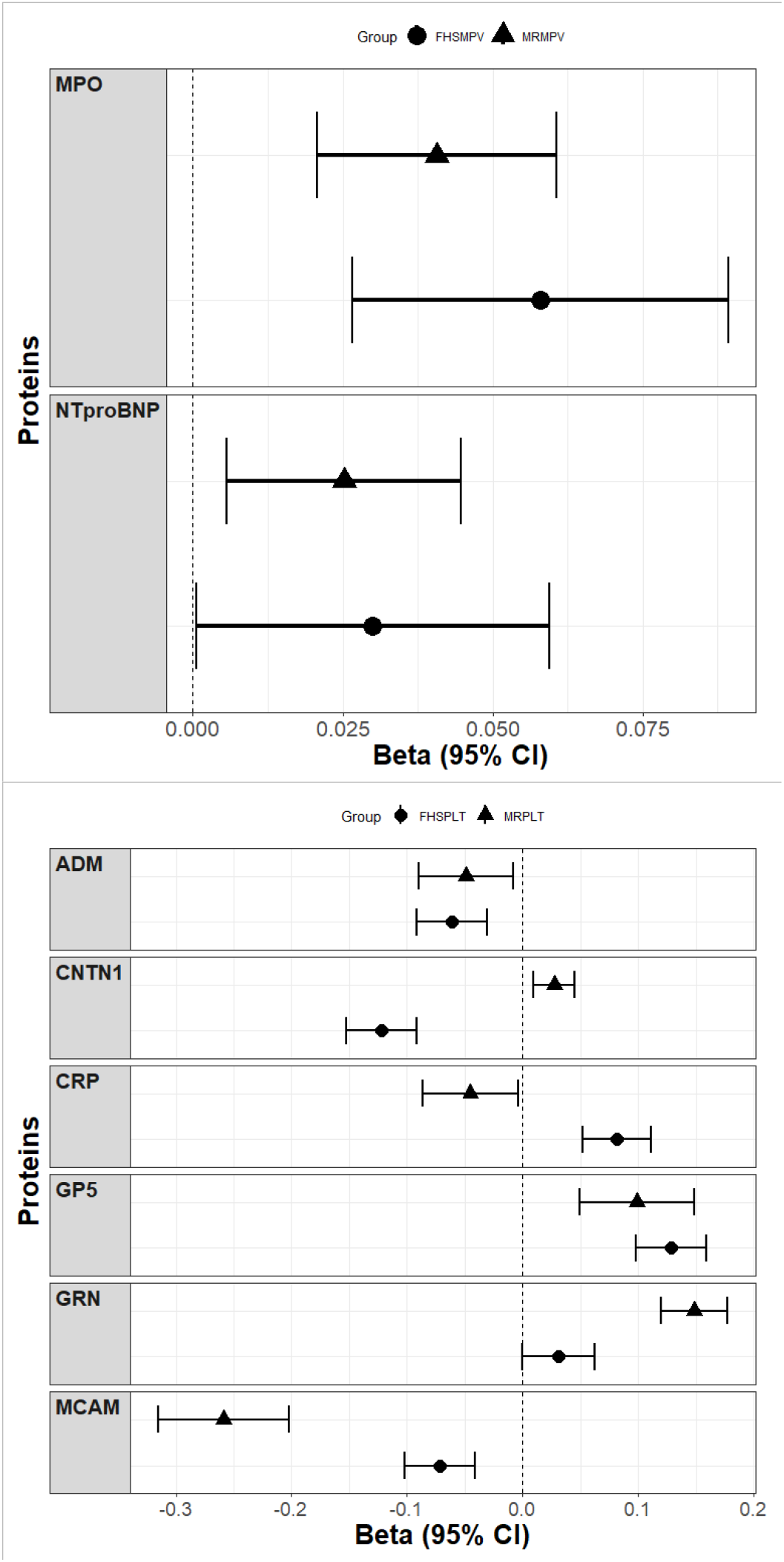
Comparison of the Effect Estimates of MR and FHS Protein-Trait Association Analyses. MPO and NTProBNP were associated with MPV (top), while ADM, CNTN1, CRP, GP5, GRN, and MCAM were associated with PLT (bottom). Proteins with P < 0.05 in both protein-trait and MR analyses were included in the plot. Units of effect estimates: per standard deviation increment of rank-based inverse normal transformed protein level.

### RNA sequence analysis of iPSC MK

To validate our findings from protein-trait association and MR analyses, we utilized human iPSC – derived MK clones that produced variable numbers of functional platelets to conduct a MK RNA-sequencing and analyzed transcriptome-wide differences between low- and high-platelet producing clones. The transcription levels of PLT-associated proteins in the iPSC MK clones showed directionally consistent associations with platelet productivity that we expected based on our human population genetics analyses (**Table 6, Supplementary Figure 4-6**).

**Table 6.** Associations Between Transcription Level and Platelet Productivity in iPSC MK Clones.

| Gene | $\beta^*$ | SE | p-value |
| --- | --- | --- | --- |
| GP5 | 0.00164 | 0.000338 | 0.00111 |
| GRN | 0.0155 | 0.00223 | 7.42E-05 |
| MCAM | -0.170 | 0.0315 | 4.34E-04 |

\*Units: platelets per megakaryocyte per fpkm

### siRNA gene knockdown experiments using iPSC MK

Motivated by our findings from the RNA sequencing analysis, we performed a siRNA gene knockdown experiment using the iPSC MK clones. Silencing GRN, GP5, and MPO genes in MK clones caused substantial reductions in platelet production (Figure 3**, Supplementary Table 6, Supplementary Figure 7**). Knockdown of MCAM appears to potentially increase platelet production, but the difference in platelet counts of MCAM knockdown and that of the control (GFP) group was not statistically significant. The gene knockdown experiments provide further support for causal association between GP5 and GRN and PLT. In addition, the experiments indicate that MPO may also be causally associated with PLT.

**Figure 3.**
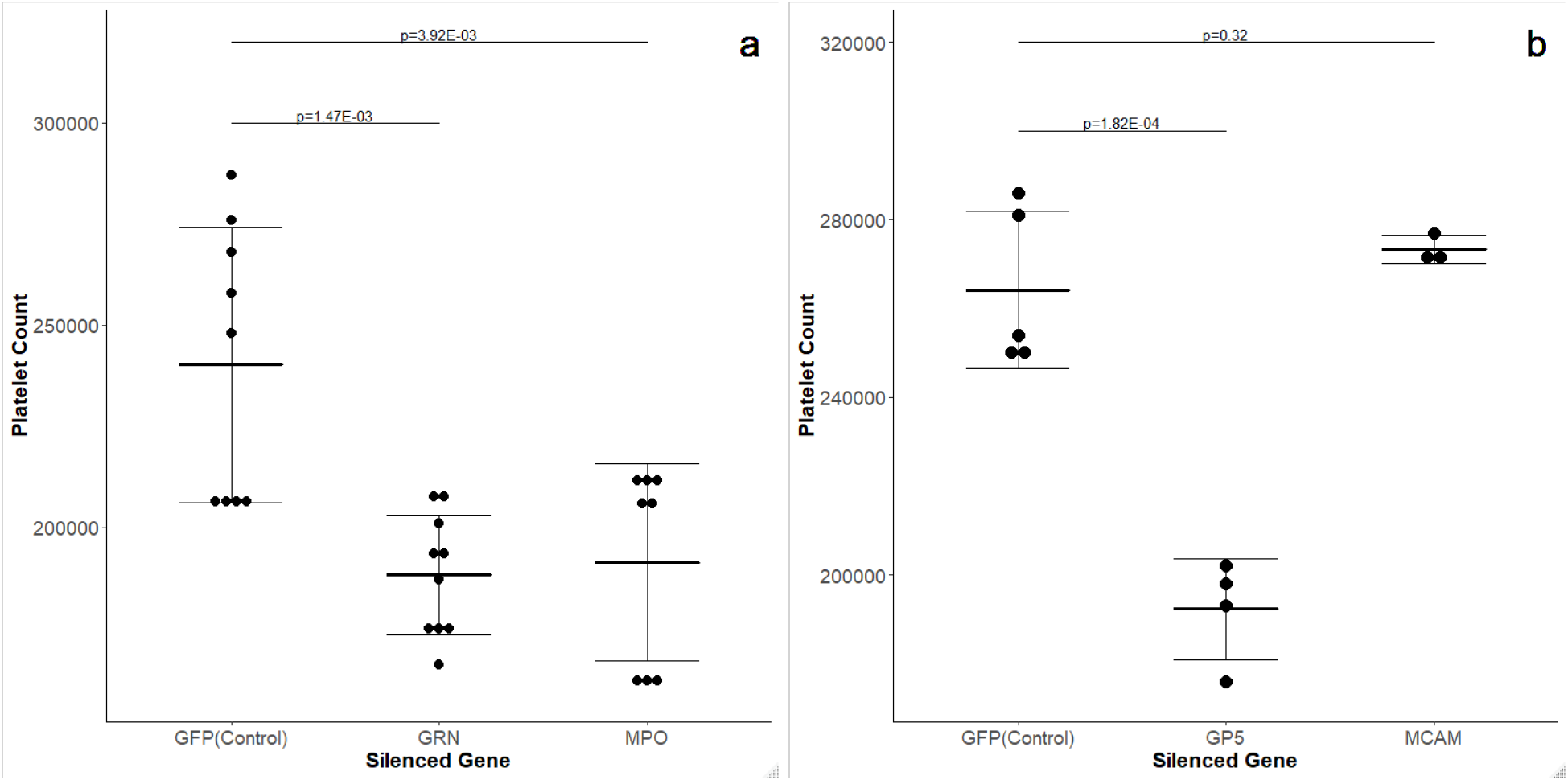
iPSC Megakaryocyte gene knockdown experiment results. **a,** Silencing GRN and MPO genes significantly decreased platelet count. **b,** Silencing GP5 significantly decreased platelet count, but silencing MCAM did not result in a significant change in platelet count.

## Discussion

There have been several large genetic studies of PLT and MPV, epidemiological studies of clinical correlates of PLT and MPV^21, 22^, and studies of plasma pQTLs^24, 37, 38^. To our knowledge, this is the first study to integrate plasma protein measurements, pQTL associations, and genetic studies using MR to identify putatively causal biomarkers of PLT and MPV followed by iPSC MK clone gene silencing experiments.

Three proteins (GP5, GRN, MCAM) were associated with PLT in FHS protein-trait, MR, as well as iPSC MK clone gene expression analyses (**Table 2**, 5**, and** 6). The causal roles of GP5 and GRN on PLT were further supported by iPSC MK gene knockdown experiments (Figure 3). GP5 is a transmembrane glycoprotein expressed on the surface of platelets. It has a soluble extracellular domain, which is cleaved by thrombin during thrombin-induced platelet activation^39^. Mutations in *GP5* are not known to cause Bernard-Soulier syndrome (enlarged platelets associated with bleeding and thrombocytopenia) unlike other gene members of the GPIb-V-IX complex. However, prior and recent work has suggested that alloantibodies targeting GP5^40^, as well as pediatric varicella^41^, gold-triggered autoimmune responses in rheumatoid arthritis^42^, and quinidine-related platelet directed antibodies^43^ may trigger immune thrombocytopenia via GP5. These studies suggest that GP5 alterations may trigger platelet clearance, and that maintenance of normal GP5 function or high levels on platelets could preserve platelet numbers in circulation. We observed consistent association between higher levels of plasma GP5 and PLT in our analyses. When we knocked down GP5 in the iPSC MK system, we observed reduced platelet production. Of note, a *trans*-pQTL SNP (rs7116008) that showed the most significant association with PLT among GP5 pQTLs (**Table 4**) is an intronic variant in gene ST3GAL4, which encodes the ST3GAL4 enzyme that is responsible for sialyation of lectin asialoglycoprotein receptor (ASGPR; Ashwell-Morell Receptor (AMR)) ligand on platelets^44^. Sialic-acid deficient platelets of ST3GAL4 knockout mice were cleared from circulation faster than platelets of wild type mice^44^. Therefore, positive correlation between GP5 and PLT might be explained by a direct effect on platelet production and an indirect effect through platelet clearance mediated by ST3GAL4 desialyation of GP5 associated with its *trans*-pQTL^44, 45^.

Granulin (GRN/epithelin) is a family of seven protein growth factors with homologous structures^46, 47^. Granulin’s precursor, progranulin (PGRN), consists of a paragranulin domain and seven GRN domains that can be cleaved and released by proteases including MMP9, MMP12, and ADAMTS-7^48^. PGRN and GRN have opposite effects on cells; GRN B promotes secretion of interleukin 8 (IL-8) by epithelial cells, but PGRN does not cause such response^46^. IL-8 has been shown to hyper-activate platelets and induce procoagulant behavior^49^. The opposing effects of GRN/PGRN on IL-8 release was also observed in human aortic smooth muscle cells^50^. PGRN appears to be anti-inflammatory^50, 51^ while GRN shows pro-inflammatory behavior^46^. PGRN has previously been implicated in blood lipid effects on CVD through its connection to sortilin (SORT1). Sortilin binds to PGRN and is responsible for endocytosis and transportation of PGRN to lysosomes^52^. A strong *trans*-pQTL for PGRN (rs646776), located on chromosome 1p13, was associated with *CELSR2* (cadherin EGF LAG seven-pass G-type receptor 2), *PSRC1* and *SORT1* (sortilin) mRNA expression^53^. SNP rs646776 was also associated with blood low-density lipoprotein (LDL) cholesterol levels and myocardial infarction risk^54^. GRN was associated with an increased risk of CVD death in a previous study^18^. Our MR analysis results using two *cis*-pQTLs of GRN (rs35203463, rs850733) and the siRNA experiment showed that silencing GRN expression leads to a decreased platelet count, suggesting a novel pathway linking GRN and CVD through platelet effects.

MPO is a heme-enzyme and a bactericidal protein found in neutrophils, specifically in azurophilic granules^55^. In addition to its role in immune response to pathogens, MPO has been suggested to play a role in inflammatory conditions and CVD^55^. MPO shows nominally significant, positive associations with coronary heart disease (CHD) events risk and CVD death in FHS data, and a positive association with CHD risk in MR^18^. In queries of publicly available GWAS data, we found consistent associations of MPO increasing alleles with CHD outcomes (**Supplementary Table 5**). MPO partially activates platelets, increasing platelets’ surface expression of P-selectin and PECAM-1 receptors as well as the frequency of aggregate formation with neutrophil granulocytes^56^. In our study, MPO was positively associated with MPV in both FHS protein-trait and MR analyses after correction for multiple testing (**Table 2** and 5). MPO also showed a positive, nominally significant (P = 0.03), association with PLT in FHS data (**Supplementary Table 2**); furthermore, MPO knockdown MK clones showed a statistically significant decrease in MK derived platelet count, demonstrating a positive correlation between the protein and platelet production (Figure 3). MPO’s positive correlation with both MPV and PLT is not concordant with the inverse relationship between MPV and PLT shown in prior literature^32^ and in our protein-trait association analysis. This difference in causal associations inferred using the MR analysis and observed in the siRNA experiment could be attributed to the tissue from which MPO was measured - MPO in FHS was measured in human plasma while the siRNA experiment measured MPO in MK cells.

Melanoma cell adhesion molecule (MCAM/CD146/MUC18) is expressed on the cell surface and contributes to binding a cell to other cells or to the extracellular matrix^57^. MCAM plays a crucial role in macrophage foam cell formation and retention of foam cells in atherosclerotic plaque^58^. Further, MCAM expression is positively correlated with necrotic core area as well as development of unstable plaques^59^. MCAM was negatively correlated with CHD risk in a prior MR study^18^. The MCAM instrumental variable SNP (rs11217234) in our study was nominally associated with (P = 0.027) with decreased risk of a second myocardial infarction event (**Supplementary Table 5**). MCAM is an important contributor to development of atherosclerosis; however, its relationship to PLT or MPV has not been studied extensively. In our study, MCAM was causally associated with decreased PLT in MR, but this inferred causality was not supported by iPSC MK gene silencing experiment (Figure 3).

Our study has several limitations. The first limitation is the time gap between plasma protein and platelet measurements in FHS participants. FHS Offspring cohort participants’ protein measurement (Exam 7) and platelet measurement (Exam 9) were about 13 years apart and FHS Third Generation cohort participants’ protein measurement (Exam 1) and platelet measurement (Exam 2) were 6 years apart. These time differences can weaken the association between protein levels and platelet traits. Second, FHS Offspring and Third Generation cohorts consist of people of European descent, potentially limiting the generalizability of study findings. The third limitation is that the proteins measured in FHS were from plasma, which might not be accurate representations of the proteins expressed in platelets.

In this study, we combined pQTL data of 71 CVD-associated proteins, platelet measurements, and published platelet GWAS to identify 4 proteins causally associated with MPV and/or PLT. MPO was causally associated with MPV and three proteins (GP5, GRN, and MCAM) were causally associated with PLT in population genomics analyses. GP5 is a well-known receptor in platelets, but MCAM and GRN are not well-characterized platelet proteins. The results of iPSC-derived MK RNA-sequencing analysis were consistent with our population-based causal associations; GRN and GP5 were over-expressed, and MCAM down-regulated, respectively, in high-platelet producing clones. Gene knockdown experiments confirmed the causal associations for GP5 and GRN in addition to suggesting that MPO might also be causally implicated in platelet production. In total, the results suggest that these proteins are causally linked to platelet generation or turnover and may play important roles in CVD via a platelet-based mechanism. Additional research on these proteins such as functional studies using mice or zebrafish may elucidate specific mechanisms through which those proteins contribute to platelet counts, potentially identifying a platelet-mediated pathway from these proteins to CVD.

## Contributions

DH.L., S-J.H., and M-H.C. conducted analyses. A.B. conducted all wet lab experiments, and A.B. and T.S. designed the imMKCL approach at Boston Children’s Hospital. DH.L. and A.D.J. wrote the manuscript. All authors critically edited the manuscript and approved the final submission.

## Acknowledgments

The views expressed in this manuscript are those of the authors and do not necessarily represent the views of the National Heart, Lung, and Blood Institute; the National Institutes of Health; or the U.S. Department of Health and Human Services.

## References

1. Machlus KR and Italiano JE, Jr. The incredible journey: From megakaryocyte development to platelet formation. J Cell Biol. 2013;201:785–796.

2. Holinstat M. Normal platelet function. Cancer metastasis reviews. 2017;36:195–198.

3. Semple JW, Italiano JE, Jr. and Freedman J. Platelets and the immune continuum. Nature reviews Immunology. 2011;11:264–74.

4. Ibrahim H and Kleiman NS. Platelet pathophysiology, pharmacology, and function in coronary artery disease. Coronary artery disease. 2017;28:614–623.

5. Davi G and Patrono C. Platelet activation and atherothrombosis. The New England journal of medicine. 2007;357:2482–94.

6. Braekkan SK, Mathiesen EB, Njolstad I, Wilsgaard T, Stormer J and Hansen JB. Mean platelet volume is a risk factor for venous thromboembolism: the Tromso Study, Tromso, Norway. Journal of thrombosis and haemostasis: JTH. 2010;8:157–62.

7. Viles-Gonzalez JF, Fuster V and Badimon JJ. Atherothrombosis: a widespread disease with unpredictable and life-threatening consequences. European heart journal. 2004;25:1197–207.

8. Sloan A, Gona P and Johnson AD. Cardiovascular correlates of platelet count and volume in the Framingham Heart Study. Annals of epidemiology. 2015;25:492–8.

9. Chu SG, Becker RC, Berger PB, Bhatt DL, Eikelboom JW, Konkle B, Mohler ER, Reilly MP and Berger JS. Mean platelet volume as a predictor of cardiovascular risk: a systematic review and meta-analysis. Journal of thrombosis and haemostasis: JTH. 2010;8:148–56.

10. Sansanayudh N, Anothaisintawee T, Muntham D, McEvoy M, Attia J and Thakkinstian A. Mean platelet volume and coronary artery disease: a systematic review and meta-analysis. International journal of cardiology. 2014;175:433–40.

11. Kim CH, Kim SJ, Lee MJ, Kwon YE, Kim YL, Park KS, Ryu HJ, Park JT, Han SH, Yoo TH, Kang SW and Oh HJ. An increase in mean platelet volume from baseline is associated with mortality in patients with severe sepsis or septic shock. PloS one. 2015;10:e0119437.

12. Gunebakmaz O, Kaya MG, Kaya EG, Ardic I, Yarlioglues M, Dogdu O, Kalay N, Akpek M, Sarli B and Ozdogru I. Mean platelet volume predicts embolic complications and prognosis in infective endocarditis. International journal of infectious diseases: IJID: official publication of the International Society for Infectious Diseases. 2010;14:e982–5.

13. Icli A, Aksoy F, Turker Y, Uysal BA, Alpay MF, Dogan A, Nar G and Varol E. Relationship Between Mean Platelet Volume and Pulmonary Embolism in Patients With Deep Vein Thrombosis. Heart, lung & circulation. 2015;24:1081–6.

14. Mayer FJ, Hoke M, Schillinger M, Minar E, Arbesu I, Koppensteiner R and Mannhalter C. Mean platelet volume predicts outcome in patients with asymptomatic carotid artery disease. European journal of clinical investigation. 2014;44:22–8.

15. Kabat GC, Kim MY, Verma AK, Manson JE, Lin J, Lessin L, Wassertheil-Smoller S and Rohan TE. Platelet count and total and cause-specific mortality in the Women’s Health Initiative. Annals of epidemiology. 2017;27:274–280.

16. Msaouel P, Lam AP, Gundabolu K, Chrysofakis G, Yu Y, Mantzaris I, Friedman E and Verma A. Abnormal platelet count is an independent predictor of mortality in the elderly and is influenced by ethnicity. Haematologica. 2014;99:930.

17. Tsai M-T, Chen Y-T, Lin C-H, Huang T-P and Tarng D-C. U-shaped mortality curve associated with platelet count among older people: a community-based cohort study. Blood. 2015;126:1633.

18. Yao C, Chen G, Song C, Keefe J, Mendelson M, Huan T, Sun BB, Laser A, Maranville JC, Wu H, Ho JE, Courchesne P, Lyass A, Larson MG, Gieger C, Graumann J, Johnson AD, Danesh J, Runz H, Hwang SJ, Liu C, Butterworth AS, Suhre K and Levy D. Genome-wide mapping of plasma protein QTLs identifies putatively causal genes and pathways for cardiovascular disease. Nature communications. 2018;9:3268.

19. Keefe J, Yao C, Hwang S-J, Courchesne P, O’Connor G, Dupuis J and Levy D. Abstract 16070: Interrogating the Proteome to Elucidate Putatively Causal Biomarkers of Emphysema: The Framingham Heart Study. 2018;138:A16070-A16070.

20. Eicher JD, Lettre G and Johnson AD. The genetics of platelet count and volume in humans. Platelets. 2018;29:125–130.

21. Eicher JD, Chami N, Kacprowski T, Nomura A, Chen MH, Yanek LR, Tajuddin SM, Schick UM, Slater AJ, Pankratz N, Polfus L, Schurmann C, Giri A, Brody JA, Lange LA, Manichaikul A, Hill WD, Pazoki R, Elliot P, Evangelou E, Tzoulaki I, Gao H, Vergnaud AC, Mathias RA, Becker DM, Becker LC, Burt A, Crosslin DR, Lyytikainen LP, Nikus K, Hernesniemi J, Kahonen M, Raitoharju E, Mononen N, Raitakari OT, Lehtimaki T, Cushman M, Zakai NA, Nickerson DA, Raffield LM, Quarells R, Willer CJ, Peloso GM, Abecasis GR, Liu DJ, Deloukas P, Samani NJ, Schunkert H, Erdmann J, Fornage M, Richard M, Tardif JC, Rioux JD, Dube MP, de Denus S, Lu Y, Bottinger EP, Loos RJ, Smith AV, Harris TB, Launer LJ, Gudnason V, Velez Edwards DR, Torstenson ES, Liu Y, Tracy RP, Rotter JI, Rich SS, Highland HM, Boerwinkle E, Li J, Lange E, Wilson JG, Mihailov E, Magi R, Hirschhorn J, Metspalu A, Esko T, Vacchi-Suzzi C, Nalls MA, Zonderman AB, Evans MK, Engstrom G, Orho-Melander M, Melander O, O’Donoghue ML, Waterworth DM, Wallentin L, White HD, Floyd JS, Bartz TM, Rice KM, Psaty BM, Starr JM, Liewald DC, Hayward C, Deary IJ, Greinacher A, Volker U, Thiele T, Volzke H, van Rooij FJ, Uitterlinden AG, Franco OH, Dehghan A, Edwards TL, Ganesh SK, Kathiresan S, Faraday N, Auer PL, Reiner AP, Lettre G and Johnson AD. Platelet-Related Variants Identified by Exomechip Meta-analysis in 157,293 Individuals. American journal of human genetics. 2016;99:40–55.

22. Astle WJ, Elding H, Jiang T, Allen D, Ruklisa D, Mann AL, Mead D, Bouman H, Riveros-Mckay F, Kostadima MA, Lambourne JJ, Sivapalaratnam S, Downes K, Kundu K, Bomba L, Berentsen K, Bradley JR, Daugherty LC, Delaneau O, Freson K, Garner SF, Grassi L, Guerrero J, Haimel M, Janssen-Megens EM, Kaan A, Kamat M, Kim B, Mandoli A, Marchini J, Martens JHA, Meacham S, Megy K, O’Connell J, Petersen R, Sharifi N, Sheard SM, Staley JR, Tuna S, van der Ent M, Walter K, Wang S-Y, Wheeler E, Wilder SP, Iotchkova V, Moore C, Sambrook J, Stunnenberg HG, Di Angelantonio E, Kaptoge S, Kuijpers TW, Carrillo-de-Santa-Pau E, Juan D, Rico D, Valencia A, Chen L, Ge B, Vasquez L, Kwan T, Garrido-Martín D, Watt S, Yang Y, Guigo R, Beck S, Paul DS, Pastinen T, Bujold D, Bourque G, Frontini M, Danesh J, Roberts DJ, Ouwehand WH, Butterworth AS and Soranzo N. The Allelic Landscape of Human Blood Cell Trait Variation and Links to Common Complex Disease. Cell. 2016;167:1415–1429.e19.

23. Mousas A, Ntritsos G, Chen M-H, Song C, Huffman JE, Tzoulaki I, Elliott P, Psaty BM, Blood-Cell C, Auer PL, Johnson AD, Evangelou E, Lettre G and Reiner AP. Rare coding variants pinpoint genes that control human hematological traits. PLOS Genetics. 2017;13:e1006925.

24. Yin X, Subramanian S, Hwang S-J, O’Donnell CJ, Fox CS, Courchesne P, Muntendam P, Gordon N, Adourian A, Juhasz P, Larson MG and Levy D. Protein biomarkers of new-onset cardiovascular disease: prospective study from the systems approach to biomarker research in cardiovascular disease initiative. Arteriosclerosis, thrombosis, and vascular biology. 2014;34:939–945.

25. Kannel WB, Feinleib M, McNamara PM, Garrison RJ and Castelli WP. An investigation of coronary heart disease in families. The Framingham offspring study. Am J Epidemiol. 1979;110:281–90.

26. Feinleib M, Kannel WB, Garrison RJ, McNamara PM and Castelli WP. The Framingham Offspring Study. Design and preliminary data. Preventive medicine. 1975;4:518–25.

27. Splansky GL, Corey D, Yang Q, Atwood LD, Cupples LA, Benjamin EJ, D’Agostino RB, Sr., Fox CS, Larson MG, Murabito JM, O’Donnell CJ, Vasan RS, Wolf PA and Levy D. The Third Generation Cohort of the National Heart, Lung, and Blood Institute’s Framingham Heart Study: design, recruitment, and initial examination. Am J Epidemiol. 2007;165:1328–35.

28. Hemani G, Zheng J, Elsworth B, Wade KH, Haberland V, Baird D, Laurin C, Burgess S, Bowden J, Langdon R, Tan VY, Yarmolinsky J, Shihab HA, Timpson NJ, Evans DM, Relton C, Martin RM, Davey Smith G, Gaunt TR and Haycock PC. The MR-Base platform supports systematic causal inference across the human phenome. eLife. 2018;7:e34408.

29. Burgess S, Small DS and Thompson SG. A review of instrumental variable estimators for Mendelian randomization. Statistical methods in medical research. 2017;26:2333–2355.

30. Haycock PC, Burgess S, Wade KH, Bowden J, Relton C and Davey Smith G. Best (but oft-forgotten) practices: the design, analysis, and interpretation of Mendelian randomization studies. The American journal of clinical nutrition. 2016;103:965–978.

31. Nakamura S, Takayama N, Hirata S, Seo H, Endo H, Ochi K, Fujita K, Koike T, Harimoto K, Dohda T, Watanabe A, Okita K, Takahashi N, Sawaguchi A, Yamanaka S, Nakauchi H, Nishimura S and Eto K. Expandable megakaryocyte cell lines enable clinically applicable generation of platelets from human induced pluripotent stem cells. Cell stem cell. 2014;14:535–48.

32. Martin-Garcia AC, Arachchillage DR, Kempny A, Alonso-Gonzalez R, Martin-Garcia A, Uebing A, Swan L, Wort SJ, Price LC, McCabe C, Sanchez PL, Dimopoulos K and Gatzoulis MA. Platelet count and mean platelet volume predict outcome in adults with Eisenmenger syndrome. Heart (British Cardiac Society). 2018;104:45–50.

33. Nelson CP, Goel A, Butterworth AS, Kanoni S, Webb TR, Marouli E, Zeng L, Ntalla I, Lai FY, Hopewell JC, Giannakopoulou O, Jiang T, Hamby SE, Di Angelantonio E, Assimes TL, Bottinger EP, Chambers JC, Clarke R, Palmer CNA, Cubbon RM, Ellinor P, Ermel R, Evangelou E, Franks PW, Grace C, Gu D, Hingorani AD, Howson JMM, Ingelsson E, Kastrati A, Kessler T, Kyriakou T, Lehtimaki T, Lu X, Lu Y, Marz W, McPherson R, Metspalu A, Pujades-Rodriguez M, Ruusalepp A, Schadt EE, Schmidt AF, Sweeting MJ, Zalloua PA, AlGhalayini K, Keavney BD, Kooner JS, Loos RJF, Patel RS, Rutter MK, Tomaszewski M, Tzoulaki I, Zeggini E, Erdmann J, Dedoussis G, Bjorkegren JLM, Schunkert H, Farrall M, Danesh J, Samani NJ, Watkins H and Deloukas P. Association analyses based on false discovery rate implicate new loci for coronary artery disease. Nature genetics. 2017;49:1385–1391.

34. Canela-Xandri O, Rawlik K and Tenesa A. An atlas of genetic associations in UK Biobank. Nature genetics. 2018;50:1593–1599.

35. Malik R, Chauhan G, Traylor M, Sargurupremraj M, Okada Y, Mishra A, Rutten-Jacobs L, Giese AK, van der Laan SW, Gretarsdottir S, Anderson CD, Chong M, Adams HHH, Ago T, Almgren P, Amouyel P, Ay H, Bartz TM, Benavente OR, Bevan S, Boncoraglio GB, Brown RD, Jr., Butterworth AS, Carrera C, Carty CL, Chasman DI, Chen WM, Cole JW, Correa A, Cotlarciuc I, Cruchaga C, Danesh J, de Bakker PIW, DeStefano AL, den Hoed M, Duan Q, Engelter ST, Falcone GJ, Gottesman RF, Grewal RP, Gudnason V, Gustafsson S, Haessler J, Harris TB, Hassan A, Havulinna AS, Heckbert SR, Holliday EG, Howard G, Hsu FC, Hyacinth HI, Ikram MA, Ingelsson E, Irvin MR, Jian X, Jimenez-Conde J, Johnson JA, Jukema JW, Kanai M, Keene KL, Kissela BM, Kleindorfer DO, Kooperberg C, Kubo M, Lange LA, Langefeld CD, Langenberg C, Launer LJ, Lee JM, Lemmens R, Leys D, Lewis CM, Lin WY, Lindgren AG, Lorentzen E, Magnusson PK, Maguire J, Manichaikul A, McArdle PF, Meschia JF, Mitchell BD, Mosley TH, Nalls MA, Ninomiya T, O’Donnell MJ, Psaty BM, Pulit SL, Rannikmae K, Reiner AP, Rexrode KM, Rice K, Rich SS, Ridker PM, Rost NS, Rothwell PM, Rotter JI, Rundek T, Sacco RL, Sakaue S, Sale MM, Salomaa V, Sapkota BR, Schmidt R, Schmidt CO, Schminke U, Sharma P, Slowik A, Sudlow CLM, Tanislav C, Tatlisumak T, Taylor KD, Thijs VNS, Thorleifsson G, Thorsteinsdottir U, Tiedt S, Trompet S, Tzourio C, van Duijn CM, Walters M, Wareham NJ, Wassertheil-Smoller S, Wilson JG, Wiggins KL, Yang Q, Yusuf S, Bis JC, Pastinen T, Ruusalepp A, Schadt EE, Koplev S, Bjorkegren JLM, Codoni V, Civelek M, Smith NL, Tregouet DA, Christophersen IE, Roselli C, Lubitz SA, Ellinor PT, Tai ES, Kooner JS, Kato N, He J, van der Harst P, Elliott P, Chambers JC, Takeuchi F, Johnson AD, Sanghera DK, Melander O, Jern C, Strbian D, Fernandez-Cadenas I, Longstreth WT, Jr., Rolfs A, Hata J, Woo D, Rosand J, Pare G, Hopewell JC, Saleheen D, Stefansson K, Worrall BB, Kittner SJ, Seshadri S, Fornage M, Markus HS, Howson JMM, Kamatani Y, Debette S and Dichgans M. Multiancestry genome-wide association study of 520,000 subjects identifies 32 loci associated with stroke and stroke subtypes. Nature genetics. 2018;50:524–537.

36. Ishigaki K, Akiyama M, Kanai M, Takahashi A, Kawakami E, Sugishita H, Sakaue S, Matoba N, Low S-K, Okada Y, Terao C, Amariuta T, Gazal S, Kochi Y, Horikoshi M, Suzuki K, Ito K, Momozawa Y, Hirata M, Matsuda K, Ikeda M, Iwata N, Ikegawa S, Kou I, Tanaka T, Nakagawa H, Suzuki A, Hirota T, Tamari M, Chayama K, Miki D, Mori M, Nagayama S, Daigo Y, Miki Y, Katagiri T, Ogawa O, Obara W, Ito H, Yoshida T, Imoto I, Takahashi T, Tanikawa C, Suzuki T, Sinozaki N, Minami S, Yamaguchi H, Asai S, Takahashi Y, Yamaji K, Takahashi K, Fujioka T, Takata R, Yanai H, Masumoto A, Koretsune Y, Kutsumi H, Higashiyama M, Murayama S, Minegishi N, Suzuki K, Tanno K, Shimizu A, Yamaji T, Iwasaki M, Sawada N, Uemura H, Tanaka K, Naito M, Sasaki M, Wakai K, Tsugane S, Yamamoto M, Yamamoto K, Murakami Y, Nakamura Y, Raychaudhuri S, Inazawa J, Yamauchi T, Kadowaki T, Kubo M and Kamatani Y. Large scale genome-wide association study in a Japanese population identified 45 novel susceptibility loci for 22 diseases. bioRxiv. 2019:795948.

37. Sun BB, Maranville JC, Peters JE, Stacey D, Staley JR, Blackshaw J, Burgess S, Jiang T, Paige E, Surendran P, Oliver-Williams C, Kamat MA, Prins BP, Wilcox SK, Zimmerman ES, Chi A, Bansal N, Spain SL, Wood AM, Morrell NW, Bradley JR, Janjic N, Roberts DJ, Ouwehand WH, Todd JA, Soranzo N, Suhre K, Paul DS, Fox CS, Plenge RM, Danesh J, Runz H and Butterworth AS. Genomic atlas of the human plasma proteome. Nature. 2018;558:73–79.

38. Ho JE, Lyass A, Courchesne P, Chen G, Liu C, Yin X, Hwang S-J, Massaro JM, Larson MG and Levy D. Protein Biomarkers of Cardiovascular Disease and Mortality in the Community. Journal of the American Heart Association. 2018;7:e008108.

39. Lanza F, Morales M, de La Salle C, Cazenave JP, Clemetson KJ, Shimomura T and Phillips DR. Cloning and characterization of the gene encoding the human platelet glycoprotein V. A member of the leucine-rich glycoprotein family cleaved during thrombin-induced platelet activation. The Journal of biological chemistry. 1993;268:20801–7.

40. Vollenberg R, Jouni R, Norris PAA, Burg-Roderfeld M, Cooper N, Rummel MJ, Bein G, Marini I, Bayat B, Burack R, Lazarus AH, Bakchoul T and Sachs UJ. Glycoprotein V is a relevant immune target in patients with immune thrombocytopenia. Haematologica. 2019;104:1237–1243.

41. Mayer JL and Beardsley DS. Varicella-associated thrombocytopenia: autoantibodies against platelet surface glycoprotein V. Pediatric research. 1996;40:615–9.

42. Garner SF, Campbell K, Metcalfe P, Keidan J, Huiskes E, Dong JF, Lopez JA and Ouwehand WH. Glycoprotein V: the predominant target antigen in gold-induced autoimmune thrombocytopenia. Blood. 2002;100:344–6.

43. Stricker RB and Shuman MA. Quinidine purpura: evidence that glycoprotein V is a target platelet antigen. Blood. 1986;67:1377–81.

44. Sorensen AL, Rumjantseva V, Nayeb-Hashemi S, Clausen H, Hartwig JH, Wandall HH and Hoffmeister KM. Role of sialic acid for platelet life span: exposure of beta-galactose results in the rapid clearance of platelets from the circulation by asialoglycoprotein receptor-expressing liver macrophages and hepatocytes. Blood. 2009;114:1645–54.

45. Jansen AJ, Josefsson EC, Rumjantseva V, Liu QP, Falet H, Bergmeier W, Cifuni SM, Sackstein R, von Andrian UH, Wagner DD, Hartwig JH and Hoffmeister KM. Desialylation accelerates platelet clearance after refrigeration and initiates GPIbalpha metalloproteinase-mediated cleavage in mice. Blood. 2012;119:1263–73.

46. Zhu J, Nathan C, Jin W, Sim D, Ashcroft GS, Wahl SM, Lacomis L, Erdjument-Bromage H, Tempst P, Wright CD and Ding A. Conversion of Proepithelin to Epithelins: Roles of SLPI and Elastase in Host Defense and Wound Repair. Cell. 2002;111:867–878.

47. Tolkatchev D, Malik S, Vinogradova A, Wang P, Chen Z, Xu P, Bennett HPJ, Bateman A and Ni F. Structure dissection of human progranulin identifies well-folded granulin/epithelin modules with unique functional activities. 2008;17:711–724.

48. Abella V, Pino J, Scotece M, Conde J, Lago F, Gonzalez-Gay MA, Mera A, Gomez R, Mobasheri A and Gualillo O. Progranulin as a biomarker and potential therapeutic agent. Drug discovery today. 2017;22:1557–1564.

49. Bester J and Pretorius E. Effects of IL-1beta, IL-6 and IL-8 on erythrocytes, platelets and clot viscoelasticity. Scientific reports. 2016;6:32188.

50. Kojima Y, Ono K, Inoue K, Takagi Y, Kikuta K-i, Nishimura M, Yoshida Y, Nakashima Y, Matsumae H, Furukawa Y, Mikuni N, Nobuyoshi M, Kimura T, Kita T and Tanaka M. Progranulin expression in advanced human atherosclerotic plaque. Atherosclerosis. 2009;206:102–108.

51. Masuda D, Nakaoka H, Komuro I, Nakatani K, Tsubakio-Yamamoto K, Nishida M, Inagaki M, Yuasa-Kawase M, Kawase R, Yamashita T, Okada T, Ohama T, Nakagawa-Toyama Y, Yamashita S, Matsuyama A, Nishihara M, Matsuwaki T and Ohmoto Y. Deletion of progranulin exacerbates atherosclerosis in ApoE knockout mice. Cardiovascular Research. 2013;100:125–133.

52. Hu F, Padukkavidana T, Vægter CB, Brady OA, Zheng Y, Mackenzie IR, Feldman HH, Nykjaer A and Strittmatter SM. Sortilin-Mediated Endocytosis Determines Levels of the Frontotemporal Dementia Protein, Progranulin. Neuron. 2010;68:654–667.

53. Kathiresan S, Melander O, Guiducci C, Surti A, Burtt NP, Rieder MJ, Cooper GM, Roos C, Voight BF, Havulinna AS, Wahlstrand B, Hedner T, Corella D, Tai ES, Ordovas JM, Berglund G, Vartiainen E, Jousilahti P, Hedblad B, Taskinen MR, Newton-Cheh C, Salomaa V, Peltonen L, Groop L, Altshuler DM and Orho-Melander M. Six new loci associated with blood low-density lipoprotein cholesterol, high-density lipoprotein cholesterol or triglycerides in humans. Nature genetics. 2008;40:189–97.

54. Carrasquillo MM, Nicholson AM, Finch N, Gibbs JR, Baker M, Rutherford NJ, Hunter TA, DeJesus-Hernandez M, Bisceglio GD, Mackenzie IR, Singleton A, Cookson MR, Crook JE, Dillman A, Hernandez D, Petersen RC, Graff-Radford NR, Younkin SG and Rademakers R. Genome-wide screen identifies rs646776 near sortilin as a regulator of progranulin levels in human plasma. American journal of human genetics. 2010;87:890–897.

55. Nussbaum C, Klinke A, Adam M, Baldus S and Sperandio M. Myeloperoxidase: a leukocyte-derived protagonist of inflammation and cardiovascular disease. Antioxidants & redox signaling. 2013;18:692–713.

56. Kolarova H, Klinke A, Kremserova S, Adam M, Pekarova M, Baldus S, Eiserich JP and Kubala L. Myeloperoxidase induces the priming of platelets. Free radical biology & medicine. 2013;61:357–69.

57. Wang Z and Yan X. CD146, a multi-functional molecule beyond adhesion. Cancer letters. 2013;330:150–62.

58. Luo Y, Duan H, Qian Y, Feng L, Wu Z, Wang F, Feng J, Yang D, Qin Z and Yan X. Macrophagic CD146 promotes foam cell formation and retention during atherosclerosis. Cell research. 2017;27:352–372.

59. Qian YN, Luo YT, Duan HX, Feng LQ, Bi Q, Wang YJ and Yan XY. Adhesion molecule CD146 and its soluble form correlate well with carotid atherosclerosis and plaque instability. CNS neuroscience & therapeutics. 2014;20:438–45.

