## Supplementary Materials for "Integrative Genomic Analysis and Functional Studies Reveal GP5, GRN, MPO and MCAM as Causal Protein Biomarkers for Platelet Traits"

**Supplementary Table 1. Oligonucleotides and siRNA used in siRNA gene silencing experiments**

| **Oligonucleotides:** |  |  |  |
| --- | --- | --- | --- |
| **qRT-PCR** | **Forward Primer (5ʹ-3ʹ)** | **Reverse Primer (5ʹ-3ʹ)** | **Amplicon size** |
| GAPDH | ACCCACTCCTCCACCTTTGA | CTGTTGCTGTAGCCAAATTCGT | 101 |
| GP5 | TTGCTTGGAGCACAGGCTAA | AGGTGGTTTCTCGAAAGCGT | 197 |
| MPO | TTCATGTTCCGCCTGGACAA | AAAAAGACCCTGCTGAGGGG | 74 |
| MCAM | CAGAAGAGATGGGCCTCCTG | GATTCGGGGCTAATGCCTCA | 98 |
| GRN | CCAGACCCACAAGCCTTGAA | TTGTCTCGGCAGCAGGTATC | 83 |
| **siRNA** |  | **Vendor** |  |
| GP5 | AUCAUGAUUUCUAAAGAAGAGCUdTdT | IDT-DNA |  |
| GFP | GCACCAUCUUCUUCAAGGAdTdT | IDT-DNA |  |
| GRN | EHU066601 | Sigma |  |
| MPO | EHU069241 | Sigma |  |
| MCAM | EHU122041 | Sigma |  |

**Supplementary Table 2. Nominally Significant (P<0.05) Protein-MPV/PLT Association Analysis Results from FHS Data**

| **Trait** | **Protein** | **β*** | **SE** | **p-value**** |
| --- | --- | --- | --- | --- |
| MPV | PMP2 | -0.049 | 0.015 | 8.48E-04 |
|  | sRAGE | 0.050 | 0.015 | 9.57E-04 |
|  | APOA1 | -0.047 | 0.014 | 1.10E-03 |
|  | KLKB1 | -0.049 | 0.015 | 1.15E-03 |
|  | BCHE | -0.042 | 0.015 | 5.31E-03 |
|  | HPX | -0.039 | 0.015 | 8.16E-03 |
|  | LDLR | -0.038 | 0.015 | 8.89E-03 |
|  | CP | -0.038 | 0.015 | 9.56E-03 |
|  | ADAM15 | -0.037 | 0.015 | 1.13E-02 |
|  | AGP (ORM) | -0.034 | 0.015 | 1.80E-02 |
|  | NRCAM | -0.034 | 0.014 | 1.94E-02 |
|  | SERPINE1 | -0.034 | 0.015 | 2.02E-02 |
|  | LEPR | -0.032 | 0.014 | 2.44E-02 |
|  | CD14 | -0.032 | 0.015 | 2.90E-02 |
|  | CD163 | -0.032 | 0.015 | 3.13E-02 |
|  | NT-proBNP | 0.030 | 0.015 | 4.60E-02 |
| PLT | CP | 0.051 | 0.015 | 8.61E-04 |
|  | LEP | 0.048 | 0.016 | 1.72E-03 |
|  | MB | -0.046 | 0.016 | 2.99E-03 |
|  | GDF15 | -0.044 | 0.017 | 7.25E-03 |
|  | MMP9 | 0.038 | 0.015 | 1.28E-02 |
|  | AMBP | 0.038 | 0.015 | 1.29E-02 |
|  | CLEC3B | -0.038 | 0.015 | 1.41E-02 |
|  | VEGF | 0.034 | 0.015 | 2.56E-02 |
|  | MCP1 | -0.033 | 0.015 | 2.75E-02 |
|  | MPO | 0.035 | 0.016 | 3.02E-02 |
|  | DPP4 | 0.033 | 0.015 | 3.38E-02 |
|  | CD14 | 0.032 | 0.015 | 3.68E-02 |
|  | GRN | 0.031 | 0.016 | 4.97E-02 |

* Units: per standard error increment in inverse-rank normalized protein level

**All proteins in the table have p-value between Bonferroni corrected significance threshold (P = 0.05/71 = 7.04E-04) and 0.05

**Supplementary Table 3. Nominally Significant (P<0.05) MR Results for MPV and PLT**

| **Trait** | **Protein** | **Number** **of** **SNPs** | **β*** | **SE** | **p-value**** |
| --- | --- | --- | --- | --- | --- |
| MPV | UCMGP | 4 | -0.029 | 0.01 | 4.16E-03 |
|  | NTproBNP | 4 | 0.025 | 0.01 | 1.14E-02 |
|  | CST3 | 11 | -0.016 | 0.0062 | 1.16E-02 |
|  | SERPINA10 | 28 | -0.007 | 0.0028 | 1.20E-02 |
|  | REG1A | 8 | -0.016 | 0.0069 | 1.79E-02 |
| PLT | CNTN1 | 7 | 0.027 | 0.0091 | 2.64E-03 |
|  | sICAM1 | 3 | 0.039 | 0.015 | 8.95E-03 |
|  | RETN | 1 | -0.069 | 0.029 | 1.76E-02 |
|  | ADM | 2 | -0.049 | 0.021 | 1.80E-02 |
|  | CRP | 2 | -0.045 | 0.021 | 3.33E-02 |

*Units: per standard error increment in inverse-rank normalized protein level

**All proteins in the table have p-value between Bonferroni corrected significance threshold (P = 0.05/37 = 1.35E-03) and 0.05

**Supplementary Table 4. SNPs Used as Instrumental Variables in MR Analysis**

| **MR Trait** | **Protein** | **SNP (RS ID)** |
| --- | --- | --- |
| MPV/PLT | GP5 | rs1466733 |
|  | GRN | rs35203463, rs850733 |
|  | SERPINA10 | rs10129500, rs1051052, rs11160183, rs116922373, rs12880244, rs138723340, rs17129523, rs186598280, rs1950843, rs20546, rs2239645, rs28520688, rs34025389, rs34878737, rs4905172, rs55951176, rs56019009, rs60461852, rs67223873, rs7143624, rs72702321, rs72706209, rs74536787, rs76249699, rs77640134, rs79768558, rs80086298, rs9972236 |
| PLT | CNTN1 | rs11178049, rs12811939, rs1838343, rs34180877, rs4768307, rs7300813, rs73110883 |
|  | MCAM | rs11217234 |
|  | CRP | rs11265266, rs2211320 |
|  | ADM | rs2218793, rs2923091 |
|  | sICAM1 | rs5498, rs74827019, rs35751322 |
|  | RETN | rs732457 |
| MPV | CD5L | rs10908610, rs2281870, rs2317232, rs2765501, rs9427315 |
|  | UCGMP | rs11056183, rs1800799, rs7135211, rs77782835 |
|  | REG1A | rs11126696, rs112649222, rs116772009, rs12471322, rs205549, rs384761, rs416188, rs76841471 |
|  | CST3 | rs113822376, rs117567509, rs117817822, rs13039514, rs17750862, rs2983289, rs4258871, rs6076118, rs76725003, rs8115833, rs911119 |
|  | NTproBNP | rs141308438, rs145790937, rs198361, rs198379 |
|  | MPO | rs2333227, rs2680703, rs34523089, rs36089182, rs79141987, rs917606 |

**Supplementary Table 5. Association of SNPs used in MR with CHD outcomes in large published GWAS**

| **Protein** | **SNP ID** | **Chr** | **Position (hg19)** | **Study** | **Trait** | **Effect**  **Allele** | **EAF** | **Effect**  **Estimate** | **p-value** | **Effect**  **Allele**  **(Yao**  **et al.)** | **β Protein** |
| --- | --- | --- | --- | --- | --- | --- | --- | --- | --- | --- | --- |
| GP5 | rs1466733 | 3 | 194120998 | UK BioBank (Canela-Xandri et al.; Roslin GeneAtlas) | Heart attack/  myocardial infarction | G | 0.237 | -0.001 | 1.94E-02 | G | -0.171 |
| MCAM | rs11217234 | 11 | 119177938 | UK BioBank (Canela-Xandri et al.; Roslin GeneAtlas) | Hypertension | G | 0.267 | 0.002 | 1.08E-02 | G | 0.144 |
|  |  |  |  | UK BioBank + CardioGramPlusC4D (Nelson et al.) | Soft CAD* | A | 0.737 | 0.023 | 1.64E-02 | A | -0.144 |
|  |  |  |  | UK BioBank (Canela-Xandri et al.; Roslin GeneAtlas) | I22 Subsequent myocardial infarction | G | 0.267 | -0.0002 | 2.71E-02 | G | 0.144 |
| GRN | rs850733 | 17 | 42451305 | UK BioBank (Canela-Xandri et al.; Roslin GeneAtlas) | Hypertension | A | 0.374 | -0.003 | 2.00E-04 | A | -0.221 |
|  |  |  |  | Japan BioBank (Ishigaki et al.; BioRxiv) | Arrhythmia | G | 0.576 | -0.032 | 6.01E-03 | G | 0.221 |
|  |  |  |  | UK BioBank (Canela-Xandri et al.; Roslin GeneAtlas) | Deep venous thrombosis (DVT) | A | 0.374 | -0.001 | 8.40E-03 | A | -0.221 |
|  |  |  |  | UK BioBank (Canela-Xandri et al.; Roslin GeneAtlas) | Venous thromboembolic disease | A | 0.374 | -0.001 | 3.66E-02 | A | -0.221 |
| GRN | rs35203463 | 17 | 42497433 | UK BioBank (Canela-Xandri et al.; Roslin GeneAtlas) | I69 Sequelae of cerebrovascular disease | C | 0.032 | 0.001 | 4.48E-03 | C | -0.362 |
|  |  |  |  | UK BioBank (Canela-Xandri et al.; Roslin GeneAtlas) | Pulmonary embolism +/- DVT | C | 0.032 | 0.001 | 1.52E-02 | C | -0.362 |
|  |  |  |  | UK BioBank (Canela-Xandri et al.; Roslin GeneAtlas) | I67 Other cerebrovascular diseases | C | 0.032 | 0.001 | 1.87E-02 | C | -0.362 |
|  |  |  |  | UK BioBank (Canela-Xandri et al.; Roslin GeneAtlas) | I60-I69 Cerebrovascular diseases | C | 0.032 | 0.002 | 2.21E-02 | C | -0.362 |
|  |  |  |  | UK BioBank (Canela-Xandri et al.; Roslin GeneAtlas) | I21 Acute myocardial infarction | C | 0.032 | 0.002 | 4.36E-02 | C | -0.362 |
| MPO | rs2333227 | 17 | 56358762 | UK BioBank (Canela-Xandri et al.; Roslin GeneAtlas) | Hypertension | T | 0.209 | 0.004 | 3.61E-05 | T | 0.146 |
|  |  |  |  | UK BioBank (Canela-Xandri et al.; Roslin GeneAtlas) | I10 Essential (primary) hypertension | T | 0.209 | 0.003 | 2.25E-03 | T | 0.146 |
|  |  |  |  | UK BioBank (Canela-Xandri et al.; Roslin GeneAtlas) | I10-I15 Hypertensive diseases | T | 0.209 | 0.003 | 2.66E-03 | T | 0.146 |
|  |  |  |  | MEGASTROKE (Malik et al.) | All stroke  (European) | T | 0.207 | 0.033 | 3.07E-03 | T | 0.146 |
|  |  |  |  | UK BioBank + CardioGramPlusC4D (Nelson et al.) | Soft CAD* | T | 0.201 | 0.028 | 6.90E-03 | T | 0.146 |
| MPO | rs2333227 | 17 | 56358762 | MEGASTROKE (Malik et al.) | Large artery stroke  (Trans-ethnic) | T | 0.201 | -0.057 | 2.78E-02 | T | 0.146 |
|  |  |  |  | UK BioBank (Canela-Xandri et al.; Roslin GeneAtlas) | I95-I99 Other and unspecified disorders of the circulatory system | T | 0.209 | 0.001 | 3.43E-02 | T | 0.146 |
| MPO | rs79141987 | 17 | 56398479 | Japan BioBank (Ishigaki et al.; BioRxiv) | Peripheral  artery disease | G | 0.710 | -0.064 | 2.15E-02 | G | 0.196 |
|  |  |  |  | UK BioBank (Canela-Xandri et al.; Roslin GeneAtlas) | Hypertension | A | 0.104 | -0.003 | 4.64E-02 | A | -0.196 |
| MPO | rs2680703 | 17 | 56429673 | UK BioBank (Canela-Xandri et al.; Roslin GeneAtlas) | Hypertension | G | 0.370 | 0.005 | 5.75E-09 | G | 0.175 |
|  |  |  |  | UK BioBank (Canela-Xandri et al.; Roslin GeneAtlas) | I10-I15 Hypertensive diseases | G | 0.370 | 0.004 | 7.88E-07 | G | 0.175 |
|  |  |  |  | UK BioBank (Canela-Xandri et al.; Roslin GeneAtlas) | I10 Essential (primary) hypertension | G | 0.370 | 0.004 | 9.03E-07 | G | 0.175 |
|  |  |  |  | MEGASTROKE (Malik et al.) | All stroke  (European) | A | 0.628 | -0.022 | 2.71E-02 | A | -0.175 |
|  |  |  |  | MEGASTROKE (Malik et al.) | All stroke,  (Trans-ethnic) | A | 0.644 | -0.020 | 3.11E-02 | A | -0.175 |
|  |  |  |  | MEGASTROKE (Malik et al.) | Cardioembolic  stroke  (trans-ethnic) | A | 0.639 | -0.040 | 3.47E-02 | A | -0.175 |
|  |  |  |  | MEGASTROKE (Malik et al.) | Cardioembolic  stroke  (European) | A | 0.628 | -0.042 | 3.70E-02 | A | -0.175 |
| MPO | rs34523089 | 17 | 56436109 | MEGASTROKE (Malik et al.) | Cardioembolic  Stroke (European) | T | 0.157 | -0.088 | 4.40E-03 | T | -0.209 |
|  |  |  |  | MEGASTROKE (Malik et al.) | Cardioembolic  stroke  (Trans-ethnic) | T | 0.155 | -0.085 | 4.81E-03 | T | -0.209 |
|  |  |  |  | MEGASTROKE (Malik et al.) | All stroke  (European) | T | 0.155 | -0.033 | 1.94E-02 | T | -0.209 |
|  |  |  |  | MEGASTROKE (Malik et al.) | All stroke  (Trans-ethnic) | T | 0.151 | -0.032 | 2.26E-02 | T | -0.209 |
|  |  |  |  | UK BioBank (Canela-Xandri et al.; Roslin GeneAtlas) | I10-I15 Hypertensive diseases | C | 0.157 | 0.002 | 3.04E-02 | C | 0.209 |

*The SOFT CAD phenotype encompasses individuals with fatal or nonfatal myocardial infarction (MI), percutaneous transluminal coronary angioplasty (PTCA) or coronary artery bypass grafting (CABG), chronic ischemic heart disease (IHD) and angina (Nelson et al).

**Supplementary Table 6. iPSC Megakaryocyte Clone siRNA Knockdown Experiment Results**

| **Group** | **Mean PLT** | **SD**  **PLT** | **Difference PLT**  **95% CI Lower** | **Difference PLT**  **95% CI Upper** | **T-test**  **p-value** |
| --- | --- | --- | --- | --- | --- |
| GFP* (control) | 240222.2 | 33984.5 |  |  |  |
| GRN | 188200.0 | 14778.4 | -79092.7 | -24951.8 | 1.47E-03 |
| MPO | 191375.0 | 24436.1 | -79317.6 | -18376.8 | 3.92E-03 |
| GFP* (control 2) | 264200.0 | 17782.0 |  |  |  |
| GP5 | 192250.0 | 11441.9 | -95262.5 | -48637.5 | 1.82E-04 |
| MCAM | 273333.3 | 3214.6 | -12712.0 | 30978.7 | 3.20E-01 |

*****Two different GFP knockdown iPSC clones were used as controls: GFP (control) was the control for siRNA knockdown experiment of GRN and MPO, and GFP (control 2) was the control for siRNA knockdown experiment of GP5 and MCAM

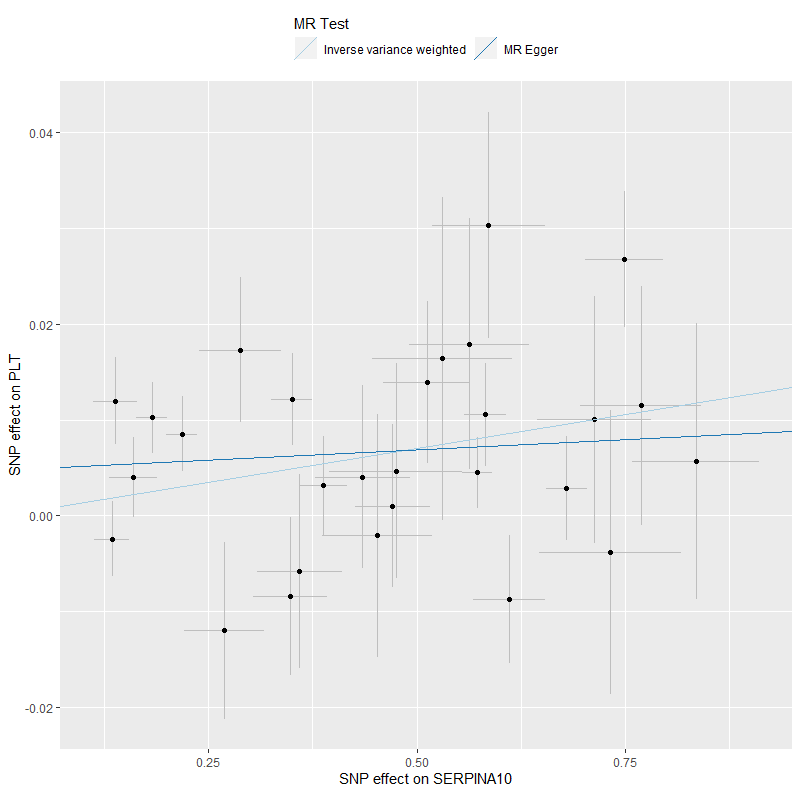

**Supplementary Figure 1. Horizontal Pleiotropy Plot for SERPINA10-PLT MR** The test for horizontal pleiotropy using 2 Sample MR package in R did not find evidence of horizontal pleiotropy among 28 cis-pQTL SNPs (egger intercept: 0.0047, se: 0.0031, p-value: 0.14)

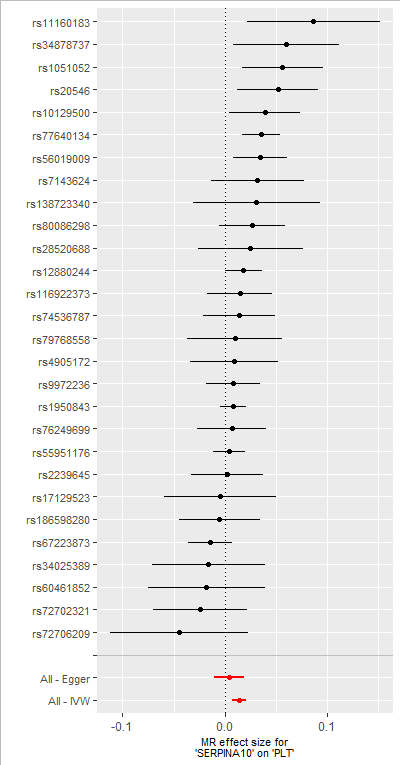

**Supplementary Figure 2. Forest Plot of MR effect size of individual cis-pQTL SNPs for SERPINA10** Forest plot revealed 7 SNPs whose MR coefficient was in the opposite direction of the overall MR direction. Those 7 SNPs are: rs17129523, rs186598280, rs67223873, rs34025389, rs60461852, rs72702321, rs72706209

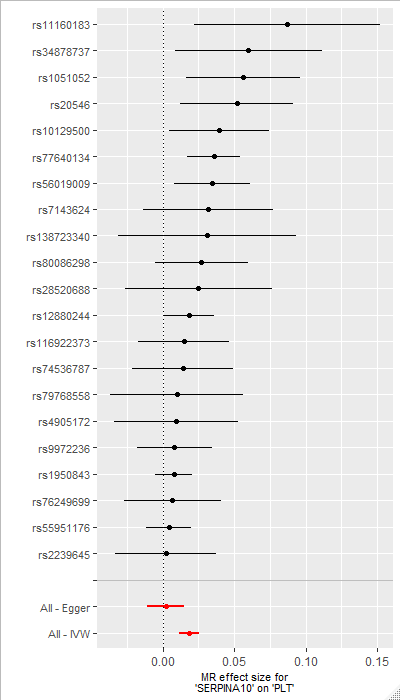

**Supplementary Figure 3. Forest Plot of MR effect size of individual cis-pQTL SNPs for SERPINA10 after removing SNPs with Negative Coefficients** The inverse-variance weighted MR coefficient using 21 SNPs was 0.019 (SE: 0.0036, p-value:1.69E-07)

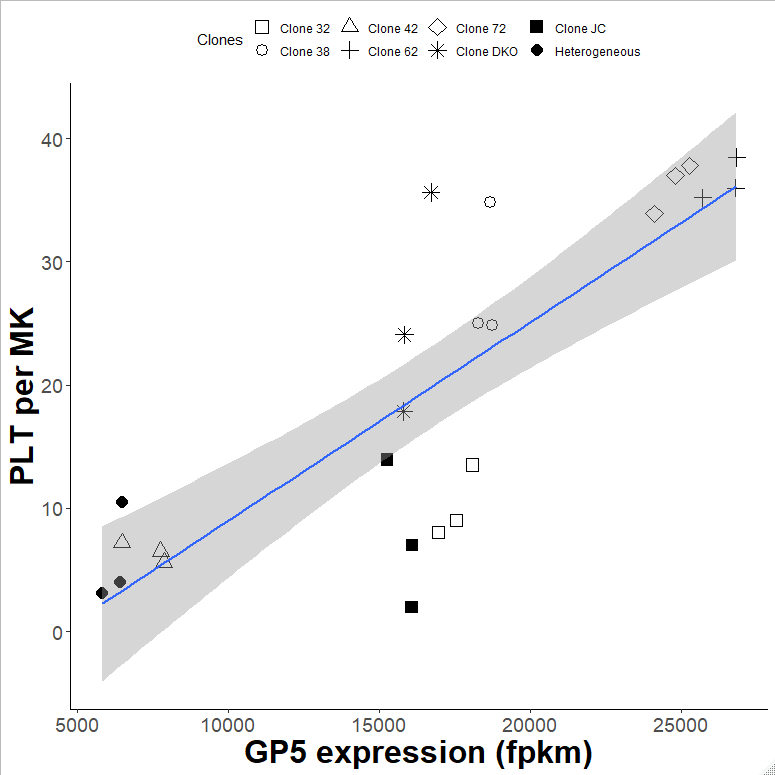

**Supplementary Figure 4. iPSC megakaryocyte GP5 transcript expression is positively associated with platelet production** Heterogenous group consists of parental cells.

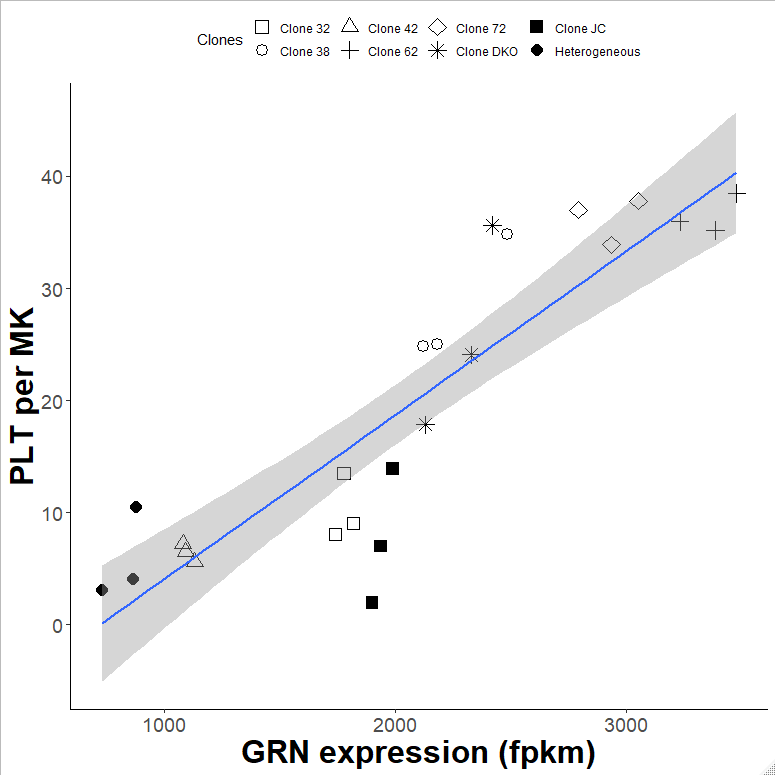

**Supplementary Figure 5. iPSC megakaryocyte GRN transcript expression is positively associated with platelet production** Heterogenous group consists of parental cells.

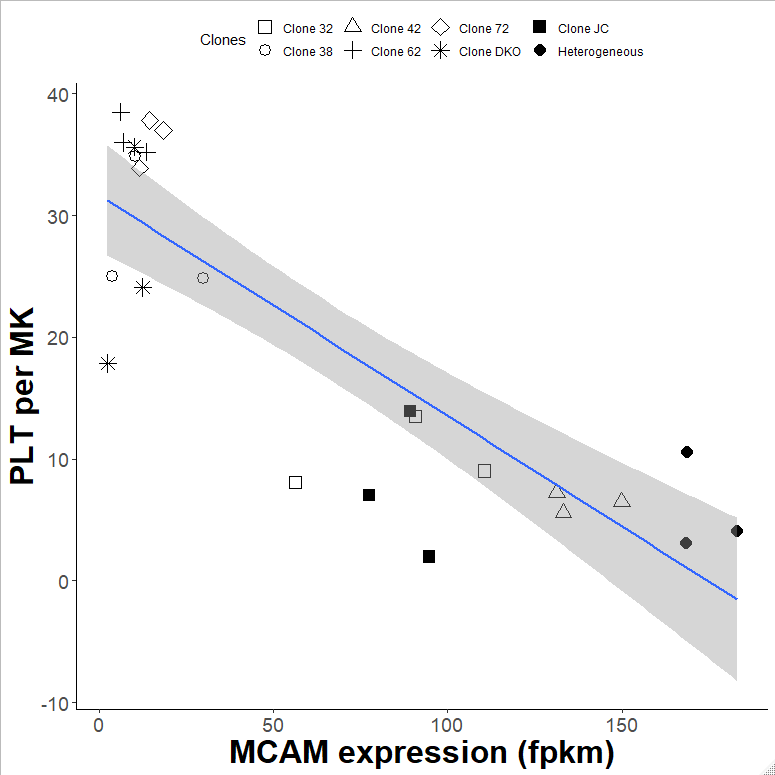

**Supplementary Figure 6. iPSC megakaryocyte MCAM transcript expression is negatively associated with platelet production** Heterogenous group consists of parental cells.

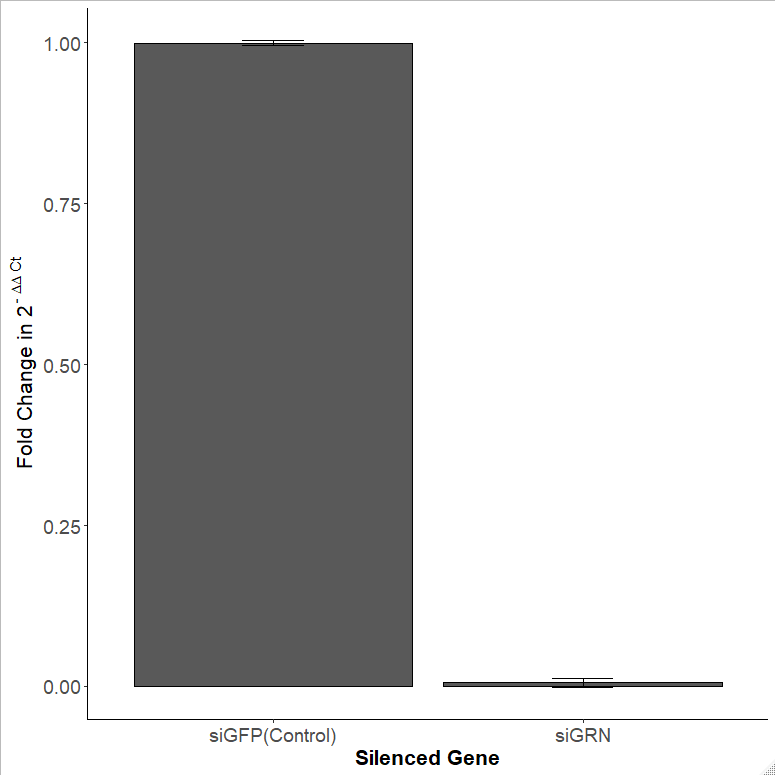

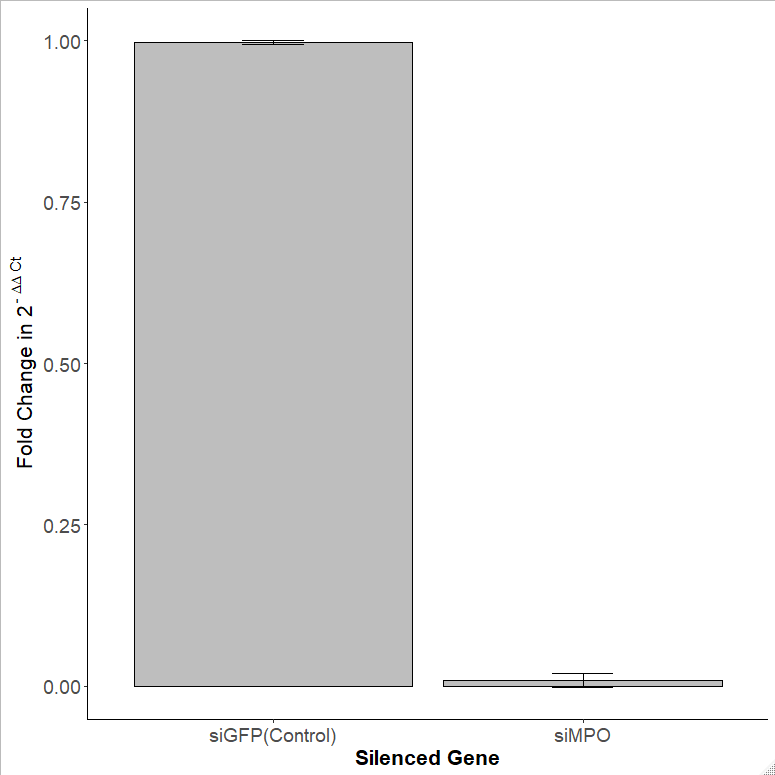

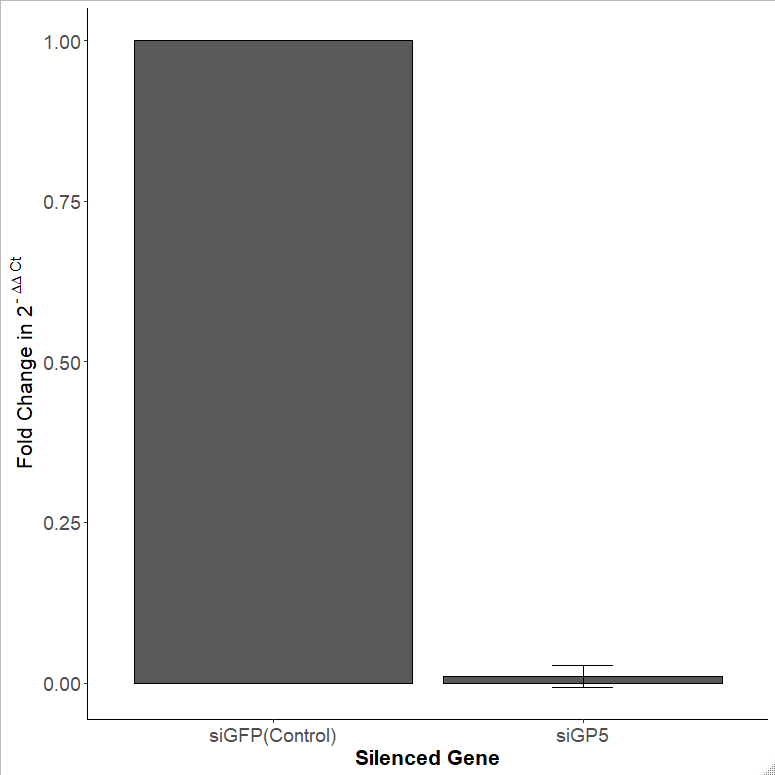

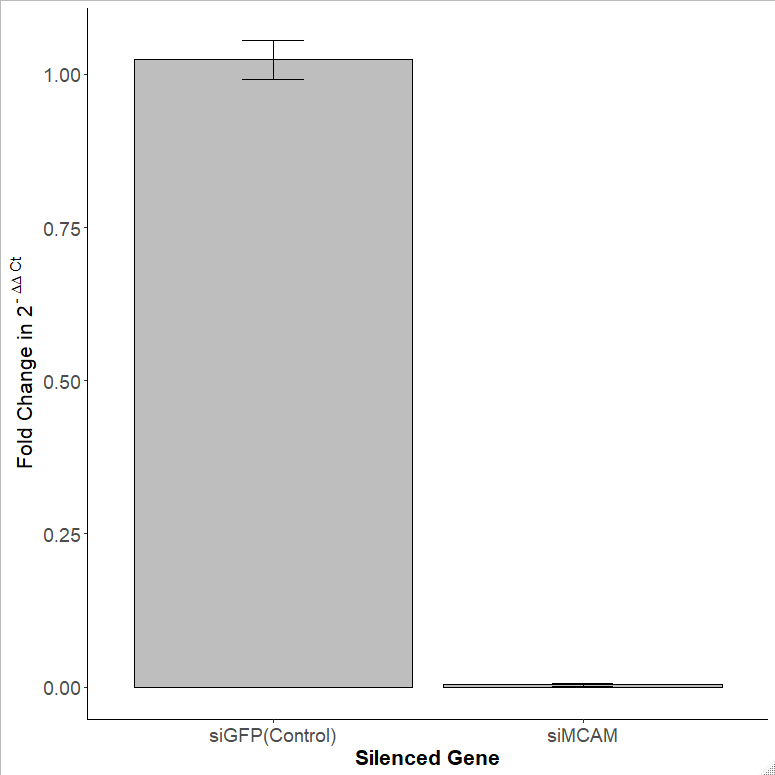

**Supplementary Figure 7. RT-qPCR plots showing decreased expression of target genes in siRNA MK clone.**
